# Supracellular organization confers directionality and mechanical potency to migrating pairs of cardiopharyngeal progenitor cells

**DOI:** 10.1101/2021.02.09.430466

**Authors:** Yelena Y. Bernadskaya, Haicen Yue, Calina Copos, Lionel Christiaen, Alex Mogilner

**Author notes:** These authors contributed equally to this manuscript (Yelena Y. Bernadskaya, Haicen Yue). These authors contributed equally to this manuscript (Lionel Christiaen, Alex Mogilner).

## Abstract

Physiological and pathological morphogenetic events involve a wide array of collective movements, suggesting that these multicellular arrangements confer biochemical and biomechanical properties that contribute to tissue scale organization. The cardiopharyngeal progenitors of the tunicate Ciona provide the simplest possible model of collective cell migration. They form cohesive bilateral cell pairs, leader-trailer polarized along the migration path as they migrate between the ventral epidermis and trunk endoderm. Here, circumventing difficulties in quantifying cellular mechanics in live embryos, we use the Cellular Potts Model to computationally probe the distributions of forces consistent with the shapes and collective polarity of migrating cell pairs. Combining computational modeling, confocal microscopy, and molecular perturbations, we first determine that cardiopharyngeal progenitors display hallmarks of supracellular organization, with differential distributions of protrusive forces, cell-matrix adhesion, and myosin-based retraction forces along the leader-trailer axis. Combined 4D simulations and experimental observations suggest that cell-cell communication helps establish a hierarchy that contributes to aligning collective polarity with the direction of migration, as observed with three or more cells both *in silico* and *in vivo*. Our approach reveals emerging properties of the migrating collective. Specifically, cell pairs are more persistent, thus migrating over longer distances, and presumably with higher accuracy. Finally, simulations suggest that polarized cell pairs literally join forces to deform the trunk endoderm, as they migrate through the extracellular space. We thus propose that the polarized supracellular organization of cardiopharyngeal progenitors confers emergent physical properties that determine mechanical interactions with their environment during morphogenesis.

## Introduction

Cell migration is a fundamental cell behavior involved in developmental and physiological processes including germline, craniofacial, and cardiac development, angiogenesis and wound healing, as well as pathogenesis such as cancer metastasis^1, 2^. In complex multicellular and dynamic environments, cells integrate biochemical and mechanical cues that guide their migration. Some migration specialists, like neutrophils, can navigate complex environments as single cells^3^. On the other hand, many developmental, homeostatic, and pathogenic morphogenetic events involve the coordinated movements of cell collectives, as observed, for example, during neural crest migration in chick, lateral line migration in zebrafish, and border cell migration in the Drosophila ovary^4 5, 6^. The properties that emerge from collective organization are thought to facilitate biochemical and mechanical integration, and thus foster efficient and accurate tissue morphogenesis in a multicellular context ^7–11^.

Migratory collectives typically exist on a continuum with varying degrees of cell-cell contacts and collective polarity, where defined leader and follower cells are arranged in a front-to-back manner^12, 13^. In minimally differentiated groups, individual cells move as autonomous units, while adjusting directionality and speed relative to their neighbors^14^. At the other extreme of collective organization, cells are integrated into supracellular arrangements, with marked front-to-back specialization and a continuity of cytoskeletal structures between neighboring cells that ensures mechanical coupling (reviewed in Shellard and Mayor, 2019). Such collective polarity implies communication between cells to coordinate subcellular processes.

Numerous studies have uncovered mechanisms underlying collective organization, such as contact inhibition of locomotion (CIL) ^13,15^ and leader-mediated inhibition of protrusive activity in follower cells^16, 17^. Ultimately, both biochemical and mechanical emerging properties of migratory collectives contribute to successful tissue morphogenesis. While many biochemical aspects of cell migration have been investigated, measurement of mechanical forces involved in *in vivo* morphogenetic cell migration has been a challenge^18^. To understand the mechanics of collective locomotion, *in vitro* techniques such as traction force microscopy were developed and used to correlate forces with movements and cytoskeletal dynamics ^19^. In more complex embryo settings, the distribution of mechanical forces can be inferred from imaging datasets, combined with available direct measurements of membrane tension^18, 20, 21^. As a complement to biophysical measurements, computational modeling offers a powerful option to reverse-engineer forces from observed cell shapes, and enable *in silico* predictions that can be compared to experimental observations^21, 22^. There is a rich inventory of modeling approaches, from simple conceptual models of cells as point-like persistent walkers interacting with distance-dependent forces ^23^, to detailed continuous or discrete models of interacting cells as distributed mechanical objects with complex rheology and free boundaries^24–26^.

We use the cardiogenic lineage of the tunicate *Ciona*, a simple chordate among the closest relatives of vertebrates^27, 28^, to develop and test a computational model of collective polarity and directed cell migration. During *Ciona* embryonic development, the cardiopharyngeal precursors migrate from their origin in the tail to the ventral trunk, hence their denomination as Trunk Ventral Cells (aka TVCs)^29–31^. The TVCs migrate as bilateral pairs, offering the simplest possible model of polarized collective cell migration. The cell in the leader position extends dynamic lamellipodia-like protrusions, while the cell in the trailer position produces a tapered retractive edge^29^. Under unperturbed conditions, the TVCs are committed to their leader/trailer positions^30, 31^. Migrating TVCs contact multiple tissues, including the posterior mesenchyme, the ventral epidermis, which serves as substrate, and the trunk endoderm^30^. During migration, the TVCs maintain polarized and bulky shapes as they invade the extracellular space between the epidermis and the endoderm. Ciona TVCs thus represent a simple and intriguing model to study the mechanics and polarity of cells migrating in an embryonic context (Fig. 1A).

**Figure 1.**
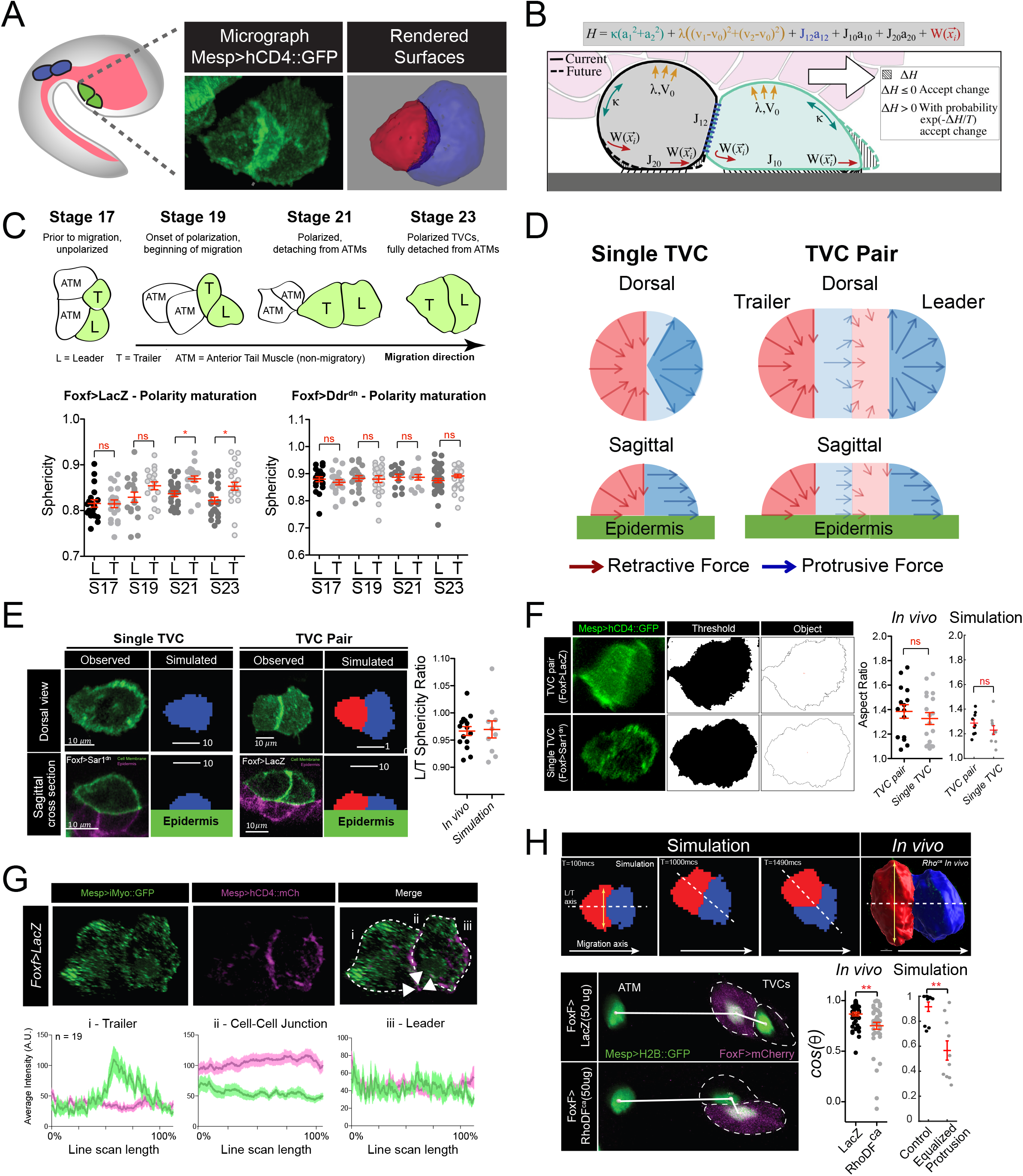
Model of force distribution in migrating TVC pairs. **(A)** Diagram of *Ciona robusta* embryo at the late tailbud stage (embryonic Stage 23). Migrating TVCs are shown in green, their non-migratory sister cells, ATMs in blue. The endoderm is shown in pink. A micrograph of a migratory pair of TVCs is shown with the leader to the right and the trailer to the left. Cell membranes are marked with Mesp>hCD4::GFP. To the right is a surface rendered image of the migratory TVC pair with leader in blue and trailer in red. **(B)** Schematic diagram showing the mechanical parameters related to cells’ movement and morphology, reflecting volume conservation (yellow), surface tension (green), cell-cell adhesion (blue), cell-epidermis adhesion (black) and active protrusion/retraction forces (red). The cell pair moves to the right, with the green cell as the leader cell and the grey cell as the trailer cell. Overlying endoderm cells are shown in pink; the underlying epidermis in grey. The shape change (shaded area) is accepted or rejected depending on the energy change Δ*H* related to it. The equation above shows the effective mechanical energy, *H*, of the cell pair. The meaning of the parameters is explained in the text. **(C)** Establishment of leader/trailer polarity as measured by the asymmetry that develops between leader and trailer sphericity as cells polarize in the direction of migration. Diagram depicts stages when sphericity was calculated. Migratory cells are highlighted in green. L = leader, T = trailer, ATM = Anterior Tail Muscle. Scatter plots show mean with standard error. Statistical significance tested using ANOVA followed by Bonferroni test to compare means. * = p<0.05 **(D)** Dorsal and sagittal views of force distributions within a single cell (left) and two connected cells (right) in our model for the unperturbed cells. Arrows indicate relative strength and direction of force. Cell anterior is in blue and posterior in red. **(E)** Comparison of cell shape in the experiment and in simulation for single migrating cell and migrating cell pair. Scatter plot shows ratio of leader to trailer sphericity derived from *in vivo* measurements and in simulations. No statistical difference was identified by Student t-test. Micrographs show dorsal and lateral view of 3D images of TVC. TVC membranes are marked with *Mesp>hCD4::GFP* and epidermal cell membranes are marked with *Mesp>hCD4::mCherry*. **(F)** Aspect ratios of migrating cell pairs compared to aspect ratios of single migrating cells calculated in FIJI and in simulations. Scatter plots show mean with standard error of *in vivo* and simulated data. Statistical analysis performed using the Student t-test. No significant difference between conditions *in vivo* and in simulation. **(G)** Distribution of myosin reporter iMyo-GFP intensity compared to membrane marker *Mesp>hCD4::mCherry*. Dashed arrows on the merged micrograph indicate the directionality of the line scan, which moves in the direction of the arrow. Mean values with standard error are plotted on the graphs. Data is derived from two pooled biological replicates. **(H)** Simulation and *in vivo* verification of equalized protrusion in leader and trailer. Top panels show results of simulated cell positions at indicated time points and the morphology of an *in vivo* cell pair when trailer protrusion is upregulated by expression of constitutively active Ras (*Ras*_*ca*_). Solid arrows show the direction of migration. Bottom panels show representative positions of migrating cells with respect to the stationary ATMs. Graphs show the cosine of the radian angle of the leader/trailer axis to the axis of migration derived from *in vivo* and simulations. Inheritance of the plasmid is followed using the cytoplasmic marker *FoxF>mCherry* (magenta) and the nuclei of the TVCs and ATMs marked with *Mesp>H2B::GFP* histone marker. *In* vivo data is pooled from two biological replicates. Statistical analysis performed using the Student t-test for the experimental data and Student t-test with Welch’s correction for the simulation data, ** = p<0.01. In simulations here and below time is measured in units of Monte Carlo Step (mcs).

Here, we model TVC shapes and behavior using the Cellular Potts Model (CPM) ^32–34^. In the CPM framework, each cell is a shifting shape described by a sum of mechanical energies of cell-substratum and cell-cell adhesions, surface tension and hydrostatic pressure, and protrusion and retraction forces (Fig. 1B). These energies effectively correspond to realistic cytoskeleton-generated forces^33 22^. The cell boundaries fluctuate, mimicking random forces and movements ubiquitous on sub-cellular scales, and shape changes minimizing the total energy are accepted, resulting in evolving, collectively moving cells (Supplemental Movies 1 and 2). While the CPM approach has some drawbacks, it is advantageous in modeling the TVC cell pair, as it allows us to reproduce both the detailed evolving 3D shapes of motile cells and the deformation of tissues surrounding these cells, over a reasonable computational time^32–34^– one of the more challenging tasks for detailed force-balance models in 3D^35^.

Here, by examining the distribution of forces required to recapitulate the shape of migrating TVCs *in silico,* we first predicted the polarized distributions of protrusive activity, cell-matrix adhesion and actomyosin contractility across cells, and tested these predictions using *in vivo* observations and molecular perturbations. We propose that the leader and trailer cells form the simplest possible supracellular arrangement of a migratory collective. We predict that this arrangement emerges from a leader-trailer mode of migration, which invokes polarized abilities to respond to extracellular guidance and mutual cell-cell attraction. Our model further explains the preference for a linear arrangement of cells polarized in the direction of migration, as this arrangement improves the persistence of migrating cells, which can presumably better buffer variations in migration cues. Finally, our model predicts that the linear arrangement of cardiopharyngeal progenitors allows them to distribute forces in a way that helps them deform the trunk endoderm and facilitates their migration despite the mechanical resistance exerted by this viscoelastic material.

## Results

### Polarized protrusive and retraction forces are distributed across a supracellular collective

Cell morphology reflects and conditions cellular behavior, inasmuch as both emerge from underlying mechanical forces (Fig. 1B) ^47^. From that standpoint, cell shapes provide a phenomenological proxy to the biophysical forces driving cellular behavior ^18, 48^. In migrating collectives, leader cells typically adopt splayed out morphologies with protrusive activity at the leading edge, while trailing cells display a tapered rear^49^. This organization is conspicuous in pairs of multipotent cardiopharyngeal progenitor cells (aka trunk ventral cells, TVCs; Fig. 1A) in the embryo of the tunicate Ciona. TVC pairs migrate along a stereotypical path, canalized by surrounding tissues, while maintaining cell-cell junctions and collectively polarizing along the direction of migration: the leader cell generates a flattened protrusive edge, while the retracting trailer cell has a tapered rear and higher sphericity^29–31^ (Fig. 1A,C,E). The Ciona TVCs thus provide the simplest possible example of directional migration of a polarized cell collective.

Maturation of TVC polarity is an active developmental process promoted by interaction of the TVCs with the extracellular matrix^31^. Taking differences in sphericity between the leader and trailer to be indicative of a fully polarized cell pair, we compared the sphericity of the leader and trailer cells at four developmental stages starting with the birth of the TVCs at developmental stage 19 prior to their migration. The TVCs’ sphericity measurements do not differ prior to migration, suggesting that they are born with equivalent shapes. As the TVCs start to move, their sphericities begin to differ significantly, indicating that they collectively polarize to adopt leader and trailer cell states (Fig. 1C) and orient them in the direction of migration. Collective TVC polarization is abolished by misexpressing a dominant negative form of the collagen receptor Ddr (Ddr^dn^), which alters integrin-mediated cell-matrix adhesion and polarized Bmp-Smad signaling^31^. This suggests that collective polarity actively matures in response to extracellular cues.

We sought to leverage the simplicity and experimental tractability of the TVC system, combined with mathematical modeling to simulate cell shape and behavior from first biophysical principles. For simplicity, we first sought to model single migrating cells, and compare simulations to observations. To this end, we use a previously established experimental perturbation, which relies on mosaic expression of a dominant-negative inhibitor of the secretory pathway (Sar1^dn^) to stall one of the TVCs, and allow the other cell to migrate on its own^30^. Single migrating TVCs generally displayed morphologies intermediate to those of either leader or trailer, generating both a leading edge and a retracting rear end, thus producing overall shapes strikingly similar to those of migrating pairs (Fig. 1E, F). Comparing the aspect ratios, defined as the ratio between the length and width of the smallest rectangular box that can enclose the cell or cell pair, of single migrating TVCs with TVC pairs shows that both maintain similar overall shapes (Fig. 1F), suggesting that similar force distributions profiles may exist in both conditions.

Cellular Potts Models or CPMs ^32–34^ and other types of models predict that cell morphologies emerge from the mechanical energies of diverse force-generating processes distributed within each cell (Fig. 1B). To explore the mechanics responsible for the observed shapes of these 1- and 2-cell systems, we selected parameters characterizing the adhesion, cortex tension and hydrostatic forces from general considerations that apply to most motile cells (see Methods), and investigated the spatial distribution of protrusive and retractive forces that would best recapitulate observed cell shapes (Fig. 1D). We first focused on the spatial-angular distribution of protrusive and retractive forces in single cells. By varying the width of the angular segment for retraction forces at the rear, we observe that narrowly focused forces cause aberrant widening of the leading edge and fail to produce the tapered rear (Supplemental Fig. 1A bottom row). This suggests that a broadly retracting trailing edge better accounts for the observed shapes of single cells.

By contrast, a wide protrusive force distribution, by extending protrusive activity to the sides, widened and flattened the leading edge (Supplemental Fig. 1A), thus better recapitulating the observed shapes (Fig. 1E). In general, several combinations of protrusive and retractive force distributions produce cell shapes that qualitatively recapitulate observations (Fig. 1E; Supplemental Fig. 1A). This suggests that the shape of motile cells is robust to variations in force distribution, while the polarized distributions of protrusive and retractile forces at the leading edge and trailing rear, respectively, is consistent with classic models of single cell migration on two dimensional substrates^50^.

Nevertheless, simulations showed that the cell shape is most faithfully reproduced if the retraction force is centripetal, radially converging to the cell center both along the width and height of the rear half of the cell, while the protrusion force is distributed radially outward along the width of the cell leading edge, but is parallel to the substrate along the height of the cell (Fig. 1D, Supplemental Movie 1). Other tested force distributions and force balance in the cell are discussed in Supplemental Information.

Empowered by our single cell simulations, we turned to modeling polarized cell pairs. Although either individual TVC can migrate on its own^30^, suggesting that individual TVCs are migration competent and do not require a cell partner, stitching together two identical cell models as defined above fails to reproduce the polarized morphology of migrating pairs. Specifically, when simulated leader and trailer cells are endowed with equivalent protrusive activity, the width of the trailer front extends beyond the rear of the leader, a morphology not observed in control embryos (Fig. 1H, yellow 2-headed arrow). This also impacts simulated migration as the equalization of forces between leader and trailer cells, through increasing protrusive activity in the trailer cell *in silico,* disrupts collective polarity causing the simulated trailer to leave its posterior position and travel more parallel to the leader (Fig. 1H). We quantified this phenomenon by calculating the cosine of the angle at which the leader/trailer axis intersects with the direction of migration (Fig. 1H). The predicted leader-dominated protrusive activity of the cell pair is consistent with previous observations that the typical leader TVC has a wide leading edge with lamellipodia-like protrusions that depend on Rhod/f- and Cdc42-controlled actin networks^29^.

Remarkably, we can reproduce this predicted behavior *in vivo*. We use the TVC-specific minimal *Foxf* enhancer to misexpress a constitutively active form of Rhod/f mosaically in either the prospective leader or trailer cell, and measure the angle between the TVCs and their stationary sister cells, the anterior tail muscles^29, 51^ (ATMs) (Fig. 1H). This assay shows that experimentally increasing protrusive activity in one of the migrating cells can disrupt their collective polarity, with cells more likely to migrate in parallel as predicted by *in silico* simulations (Fig. 1H). This result suggests that the protrusive activity is suppressed at the anterior of the trailer cell compared to leader and single cells.

However, some protrusive activity in the trailer may still be required, as numerical simulations that reduce protrusive activity in the trailer without increasing its retractile forces cause the leader to detach and move forward on its own overcoming significant mutual adhesion between the two cells (Supplemental Fig. 1C). This suggests that in paired TVC migration cells exert roughly equivalent forces and coordinate their activities to distribute protrusive and retractile forces to the leader and trailer cells, respectively. We discuss all possible force combinations in two adhesive cells in the Supplemental section (Supplemental Fig. 1B-E).

Next, we sought to further probe the supracellular model using phenomenological observations and experimental perturbations. Notably, the best computational recapitulation of observed shapes was achieved with centripetal retractive forces dominating in the trailer, pulling the cell rear forward and down, while the protrusive forces in the leader pushed the front forward parallel to the substrate (Fig. 1D, Supplemental Movie 2). Additionally, small protrusive forces in the leader and retractive forces in the trailer assist the dominant protrusive/retractive forces in the leader/trailer, respectively, as described above. This force distribution also reproduced the measured aspect ratio of the motile TVC pair and single cells (Fig. 1F) and the morphological asymmetries of the leader and trailer as reported by the ratio of their sphericities in vivo compared to simulations (Fig. 1E). This is reminiscent of the centripetal character of actomyosin contractility ^52^ with planar alignment of protrusive forces within layered lamellipodial actin arrays^53^. Consistent with the model’s prediction of a trailer-polarized retractile activity, we observe that iMyo-GFP, an intrabody that recognizes non-muscle myosin II through a conserved epitope^38, 54^ accumulates at the rear of the trailer cell, and is relatively depleted from the leader-trailer junction (Fig. 1G), implying that the retraction in the trailer is dominant, while the retractive force in the leader is weak, further supporting our interpretation of the TVC pair as a supracellular structure.

Taken together, our computational predictions and experimental observations support a model where pairs of multipotent cardiopharyngeal progenitors migrate as a polarized supracellular collective, with protrusive activity and myosin-based retraction distributed across leader and trailer cells, respectively.

### Polarized cell-matrix adhesion contributes to leader/trailer state of migrating cells

We previously showed that Integrin ß1(Intß1)- and Discoidin domain receptor (Ddr)-mediated signaling and cell-matrix adhesion to the basal lamina of the ventral trunk epidermis promote collective polarity and directional movement of TVC pairs^31^ (Fig. 1C). Here, we harness the predictive power of our model to explore the phenotypic consequences of numerically varying the distribution and strength of cell-matrix adhesion forces across the cell pair *in silico*. We begin by simulating two cells side-by-side with the same protrusive/retractive forces distributions, but with cell-matrix adhesion in one of the cells lower than the other. Under these conditions we find that the cell with reduced adhesion is more likely to assume the trailer position (Fig. 2A). We tested this prediction *in vivo* using mosaic overexpression of the dominant negative form of Integrin ß1 driven by the *Foxf* minimal TVC enhancer (Fig. 2B). In control mosaic embryos, *Foxf*_*TVC*_-driven fluorescent protein expression marks either the leader or trailer cell in equal proportions, consistent with previously published data^30^ (Fig. 2B). By contrast, co-expression of *Intß1*^*dn*^ increases the proportion of labeled trailer cells to 57% in mosaic embryos (Fig. 2B). Further, one simulation of a more acute reduction of cell*-*matrix adhesion in the trailer of leader-trailer polarized cell pairs (see Sup Table) resulted in a rare *in silico* tumbling behavior (Fig. 2C), where the low-adhesion trailer cell climbs on top of and over the normally adhering leader cell. Notably, we previously observed this distinct cell behavior *in vivo* (experimental image is shown in Fig. 2C), following TVC-specific inhibition of cell-matrix adhesion, and reduction of collagen9-a1 secretion from the adjacent trunk endoderm^30, 31^. Taken together, these observations indicate that reduced cell-matrix adhesion promotes positioning of the cell with reduced adhesion posterior to the cell with the larger adhesion, thereby preferentially adopting the trailer state, which in turn suggests that cell-matrix adhesion is stronger in leader cells.

**Figure 2.**
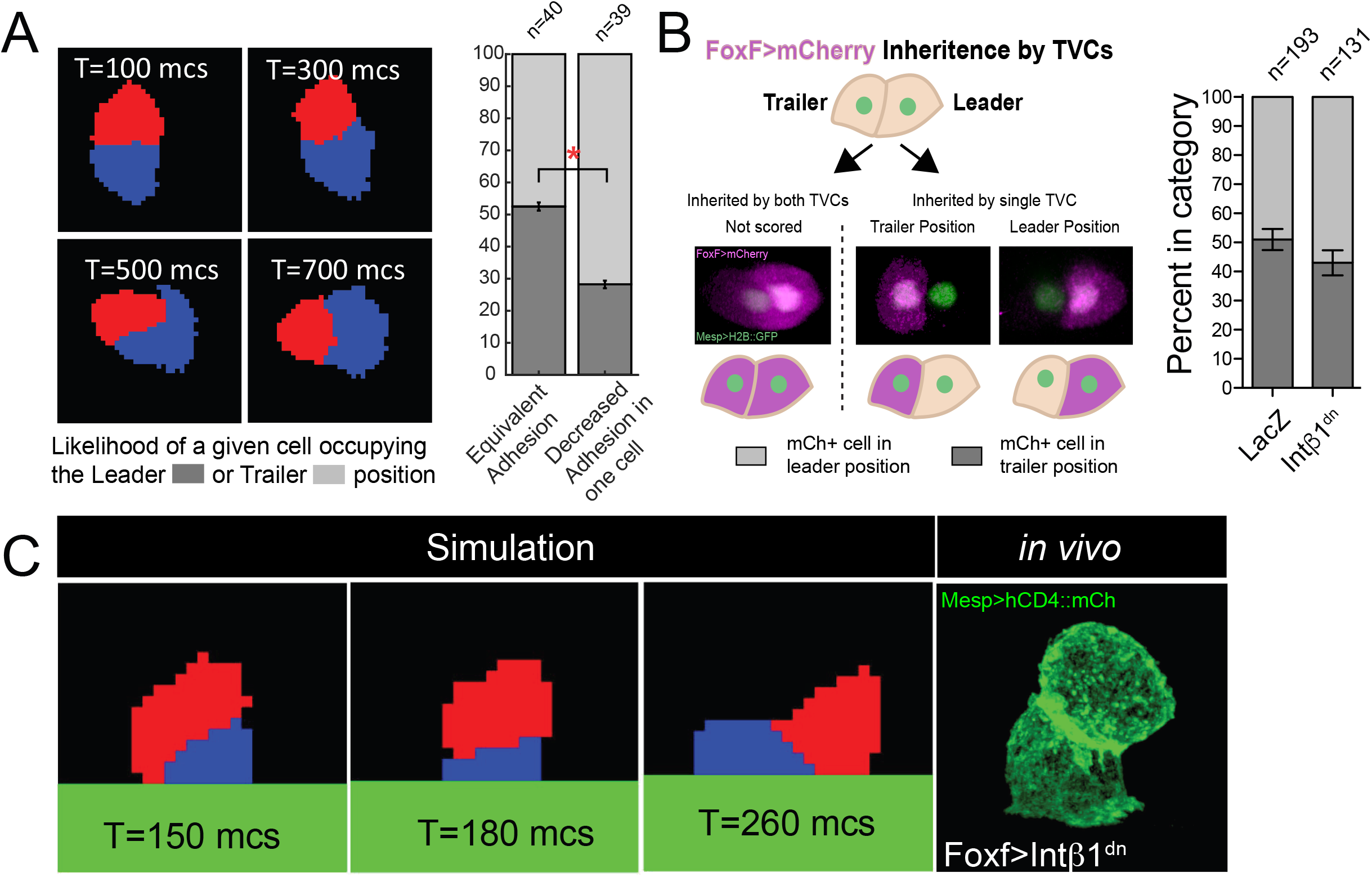
Polarized matrix adhesion promotes adoption of leader/trailer cell state. **(A)** Simulation of decreasing ECM adhesion in one cell (red) of a migrating cell pair. Cells are migrating to the right starting in a parallel orientation as shown in T=100 mcs (Monte Carlo Steps). Bar graphs show likelihood of either cell assuming the leader or trailer position in either control conditions (50/50 likelihood) or when adhesion in red cell is decreased. Standard error of proportion is shown and statistical analysis of the proportions is done using Fisher’s exact test. **(B)** Invivo modulation of ECM adhesion using mosaic inheritance of the *Foxf>Intb1*^*dn*^ marked by *Foxf>mCherry*. Diagram shows a schematic of mosaic inheritance of transgenes and resulting distribution of mCherry fluorescence. Bar graph shows likelihood of cell that inherits the transgenic constrict to be found in either leader or trailer position. Error bars are standard error of proportion. Statistical analysis using Fisher’s exact test does not find statistical significance between conditions. **(C)** Acute reduction of ECM adhesion in single cell (red) causes detachment of that cell from the underlying epidermis and recapitulates the phenotype observed in vivo (bottom right) with the detached cell positioned on top of the cell that maintains ECM adhesion. Micrograph at the bottom right shows TVC pair expressing *Foxf>dnIntb1*^*dn*^ with membranes marked by *Mesp>hCD4::GFP*.

### Hierarchical guidance orients collective polarity in the direction of migration

From the above sections, a picture emerges whereby the supracellular organization of migrating pairs of cardiopharyngeal progenitors is characterized by leader-polarized protrusive activity and cell-matrix adhesion, and trailer-polarized deadhesion and myosin-driven retraction. Both experimental and simulated disruptions of this supracellular polarity alter directionality, which is marked by an alignment of the leader-trailer axis with the direction of migration. However, the two cells are not arranged in a linear leader-trailer orientation at birth (Fig. 3A,G,), and previous observations indicated that either cell can assume a leader position, although single cell lineage tracing indicated that the leader emerges from the most anterior founder cell in ~95% of the embryos^30^.

**Figure 3.**
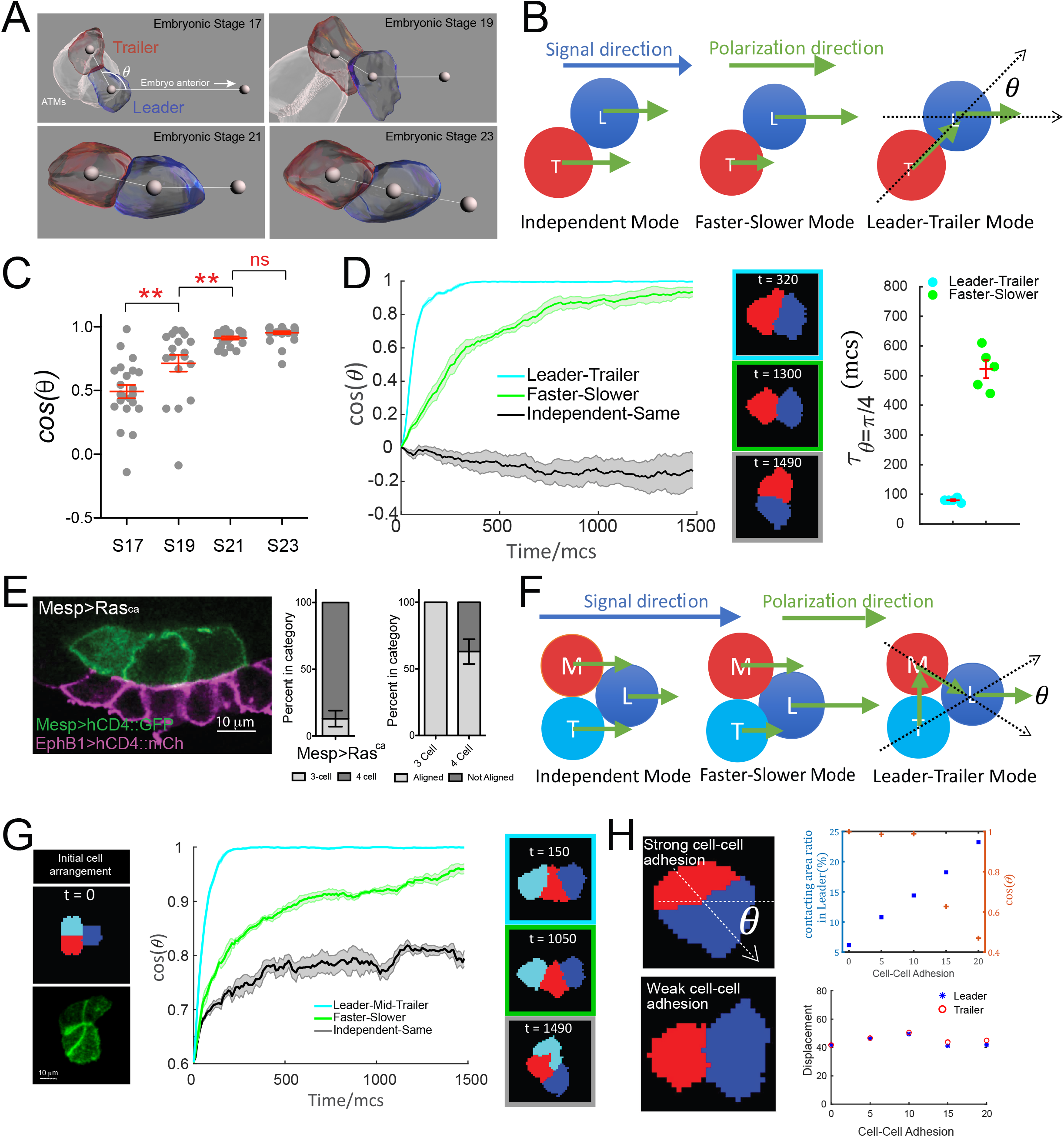
Hierarchical organization of multicellular migratory clusters. **(A)** Evolution of TVC polarization. Panels show in vivo rendered images of cells at the indicated embryonic stages. Leader in blue, trailer in red, non-migratory ATMs in white. Spheres inside cells mark the center of mass, sphere to the right indicates direction of anterior migration. Angle theta between the axis of leader/trailer and direction of migration is indicated. **(B)** Three hypothesized polarization modes for two cells. Independent: cells polarize independently in the signal direction and move with the same speed. Faster-Slower: cells polarize independently in the signal direction, but one cell moves faster than the other. Leader-Trailer: one cell (leader) follows the signal direction, while the other (trailer) polarizes in the direction of the leader’s center-of-mass. L = leader, T = trailer. **(C)** Establishment of alignment between the leader/trailer axis and direction of migration. *Cos(q)* is shown for indicated embryonic stages. Data is pooled from two biological replicates. Statistical analysis performed using 1-way ANOVA and Bonferroni post test. ** = p<0.01. **(D)** Left: The simulated evolution of two cells’ geometry (quantified as cosine of the angle between the line connecting the cells centroids and the signal direction, *(COSθ*) for the three polarization modes shown in (B). 5 simulations are run for each mode and the shaded area shows the standard error. Center: Representative snapshots when the two cells reach linear arrangement or at the end of the simulation using each mode three modes. The colors of the frames correspond to the data set on the graph. Right: Scatter plot showing the time when *θ* reaches π/4 for two modes with mean and standard error and statistical analysis using student t-test with Welch’s correction. **(E)** Three migratory cells are linearly arranged in the direction of migration. Bar graphs show the proportion of TVCs that migrate as either 3 or 4 cells under induced MAPK signaling by *Mesp>Ras*^*ca*^ and proportion of cell groups that are linearly polarized in each subset. Error bars show standard error of proportion. **(F)** Hypothesized polarization modes for three cells. Independent: cells polarize independently in the signal direction and move with equal speeds. Faster-Slower: cells polarize independently in the signal direction, in this case the leader travels the fastest, trailer the slowest and middle cell travels at an intermediate speed. Leader-Trailer: one cell (leader) follows the signal direction, middle cell polarizes towards the leader and trailer polarizes toward the middle cell. **(G)** Simulation of the 3-cell group polarization under the three hypothesized polarization modes. Left: the initial cell arrangement *in silico* (top) and *in vivo* (bottom). Note that *in vivo* there are always four cells prior to migration. Center: the polarization of migrating cell clusters over time is quantified by the cosine of the angle between the lines connecting the leader and the two posterior cells separately, as shown in (F). 5 simulations were run for each mode and shaded area shows the standard error. Right: representative snapshots when the three cells reach linear arrangement for the three modes examined (for the Independent-Same mode linear arrangement is never reached, snapshot shows cells at the end of a simulation run). The colors of the frames correspond to the data sets on the graph. **(H)** Effectsof modulating cell-cell adhesion on the contacting area between the two cells and on their speed, quantified by the percentage of total surface area of the leader cell (top graph, left y-axis, blue symbols), on the ability of the cell pair to polarize in the direction of migration quantified by *COSθ* (top graph, right y-axis, red symbols), where *θ* is the angle between the line connecting two cells and the moving direction as shown in the top image on the left, and on the total displacement of the leader/trailer pair (bottom graph). x-axis shows the relative energy of the cell-cell junction (the adhesion parameter is rescaled here so that larger value means stronger cell-cell adhesion). Images show cell pairs with either high (top) or low (bottom) cell-cell adhesion. Arrow represents leader/trailer axis.

We analyzed the establishment of leader/trailer polarity from the initial time of TVC birth to full polarization and alignment with the direction of migration over 4 embryonic stages that encompass TVC migration^55^. We use *cos(θ)* to quantify the alignment of cell pairs with the direction of migration, with *θ* defined as the angle between the leader-trailer axis (axis connecting their centers-of-mass) and the direction of migration (Fig. 3A,B). Prior to migration at embryonic stage 17 (8.5 hours post fertilization, hpf), the leader trailer axis is more orthogonal to the direction of movement. The cells reach their full polarization and alignment with direction of migration by stage 21, a process that takes approximately 1.5 hours at 18 °C, after which they continue to migrate as a fully polarized cell pair for approximately two hours until stage 23 (Fig. 3A,C).

We thus sought to explore possible mechanisms governing the establishment of collective leader-trailer polarity and its alignment with the migration path. We modeled three possible directional modes for the cell pair (Fig. 3B): the Independent Mode, where the two cells have the same distributions of the retractive and protrusion forces and their polarization directions – retractive-protrusive axes – are both aligned to the right, along the net directional signal from the surrounding tissue; the Faster-Slower Mode, where the total retractive-protrusive force in one cell is scaled up compared to that in the other cell, thus making one cell faster than the other; and the Leader-Trailer Mode, where the prospective leader’s retractive-protrusive axis is aligned with the external signal direction, while the prospective trailer’s retractive-protrusive axis is oriented toward the leader’s center-of-mass. In all these modes, the directional noise is not included. In simulations, the two cells are initially placed side-by-side, with their retractive-protrusive axes orthogonal to the migration path (i.e. *cos(θ) = 0*), thus resembling the arrangement observed *in vivo* prior to migration onset. Simulations show that the cells following the Independent mode maintain their side-by-side orientation, and fail to align with the migration path (Fig. 3D, Supplemental Movie 3,4). By contrast, either the Faster-Slower or Leader-Trailer mode allows the cells to rearrange into a single file aligned with the migration path (i.e. *cos(θ) = 1*), with the Leader-Trailer mode allowing alignment to occur earlier, with a half-time to alignment reduced compared to the Fast-Slower mode (Fig. 3D). This predicted behavior agrees qualitatively with the observed polarization of TVCs *in vivo* described above.

The assumption that basic rules of polarization of follower cells towards the cells anterior to them governs the hierarchical adoption of leader and trailer states predicts that the linear arrangement of cells should be maintained even if the number of migrating cells were increased. Indeed, we previously observed that ectopic FGF/M-Ras/Mek-driven induction within the *Mesp*+ lineage causes 3 or 4 cells to assume a cardiopharyngeal identity and migrate collectively^29, 56, 57^. In conditions such as misexpression of a constitutively active form of M-Ras using the *Mesp* enhancer^57^ (*Mesp>M-Ras*^*ca*^), the cells align in the direction of movement in 57% of the experimental embryos, with a single anterior leader, followed by 2 (13%) or 3 (87%) cells arranged in a single line (68%, n = 31) (Fig. 3E; Supplemental movie 5, 6).

To test this, we further probed the three distinct migration modes by modeling three migrating cells (Fig. 3F). In these simulations, each of the three cells adheres equally to the other two. The modes are similar to those for two cells (Fig. 3B), with two adaptations: in the Faster-Slower mode, one cell is the fastest, another – the slowest, and the third one moves with an intermediate speed; the Leader-Trailer mode becomes the Leader-Middle-Trailer mode, in which the polarization axis of the leader is fixed to the external signal direction, the middle cell’s axis orients toward the leader’s center, and the trailer’s axis orients toward the middle cell’s center. To mirror the initial arrangement observed *in vivo*, we start 3-cell simulations with individual cells distributed in an approximately triangular pattern (Fig. 3G, Supplemental Movie 7,8). Similar to two-cell simulations, the Independent Mode failed to produce linear arrangements of cells. Under these parameters, the cells remained in the triangular formation with no specific leader emerging. This is likely due to the 3-cell system minimizing the adhesive energy when each cell maintains contacts with the other two (Fig. 3G). Although cells arranged more linearly under the Faster-Slower mode (Fig. 3E, Supplemental Movie 7), they failed to align with the direction of migration for an extended period of time, thus only achieving the single-file arrangement toward the end of simulations (Fig. 3G). The Leader-Middle-Trailer mode was again the most effective at producing full polarization and linear order rapidly (Fig. 3G, Supplemental Movie 8), thus suggesting that basic rules of collective polarization can produce hierarchical arrangements of cell groups containing variable numbers of cells.

Since the ability to organize cell collective requires maintenance of cell-cell junctions, we also investigated the effect of the cell-cell adhesion strength on collective polarization. To this end, we performed multiple simulations by varying the cell-cell adhesion energy in our basic model. Simulations showed that cell-cell adhesion strength does not affect the total displacement of cell pairs over long time intervals (Fig. 3H, bottom right). However, increasing/decreasing cell-cell adhesion energy caused the cell-cell boundary area to increase or decrease, respectively (Fig. 3H, top right, blue plot), which leads to a drastic misalignment of cells with the direction of migration above a certain threshold. This suggests that the extent of cell-cell adhesion may be regulated *in vivo* and that highly adhesive cells will reorient their supracellular polarity away from the direction of migration.

Taken together, the above analyses indicate that when two or more cells move together, they establish a hierarchical collective polarity, whereby a combination of extrinsic guidance, leader-trailer order, as well as possibly differential force generation and/or regulation of cell-cell adhesion strength cause a single leader to emerge, followed by one or more cells that organize and maintain a single file aligned with the direction of migration.

### Collective polarity fosters persistent directionality

Having characterized ground rules that govern the collective arrangement of migrating cardiopharyngeal progenitors, we sought to explore the specific properties conferred by this supracellular organization. Unlike specialized motile cells, such as Dictyostelium, fish keratocytes or neutrophils, which can migrate at ~ 10 μm/min, 7.5 μm/min, ~ 19 μm/min, respectively^58–60^, TVCs move at ~ 0.4 μm/min, which is relatively slow, but not unexpected for a developmental migration that contributes to the establishment of accurate cellular patterns in the embryo^61^. We reasoned that this behavioral accuracy would be reflected in the cells’ persistence, defined as the ratio of beginning-to-end displacement to trajectory length^62^ (Fig. 4A). Countering persistence and accuracy, directional noise, which can emerge from inherent stochasticity of motile engines, from random fluctuations of external cues, and/or signal transduction^63^, can cause meandering trajectories of cells^64^. We therefore simulated noise by adding directional stochasticity to the model of two cells in the Leader-Trailer mode, and compare the persistence of migrating cell pairs with that of single cells (Fig. 4A, B). In the simulations, motile cell pairs were always more persistent than single cells when centers of mass were tracked over time for each condition, suggesting that single cells are more sensitive to directional stochasticity. Interestingly, the total length of the migration path in the simulations was not altered, suggesting that the decreased displacement is a function of the meandering path traveled by the less persistent single cells (Figure 4B).

**Figure 4.**
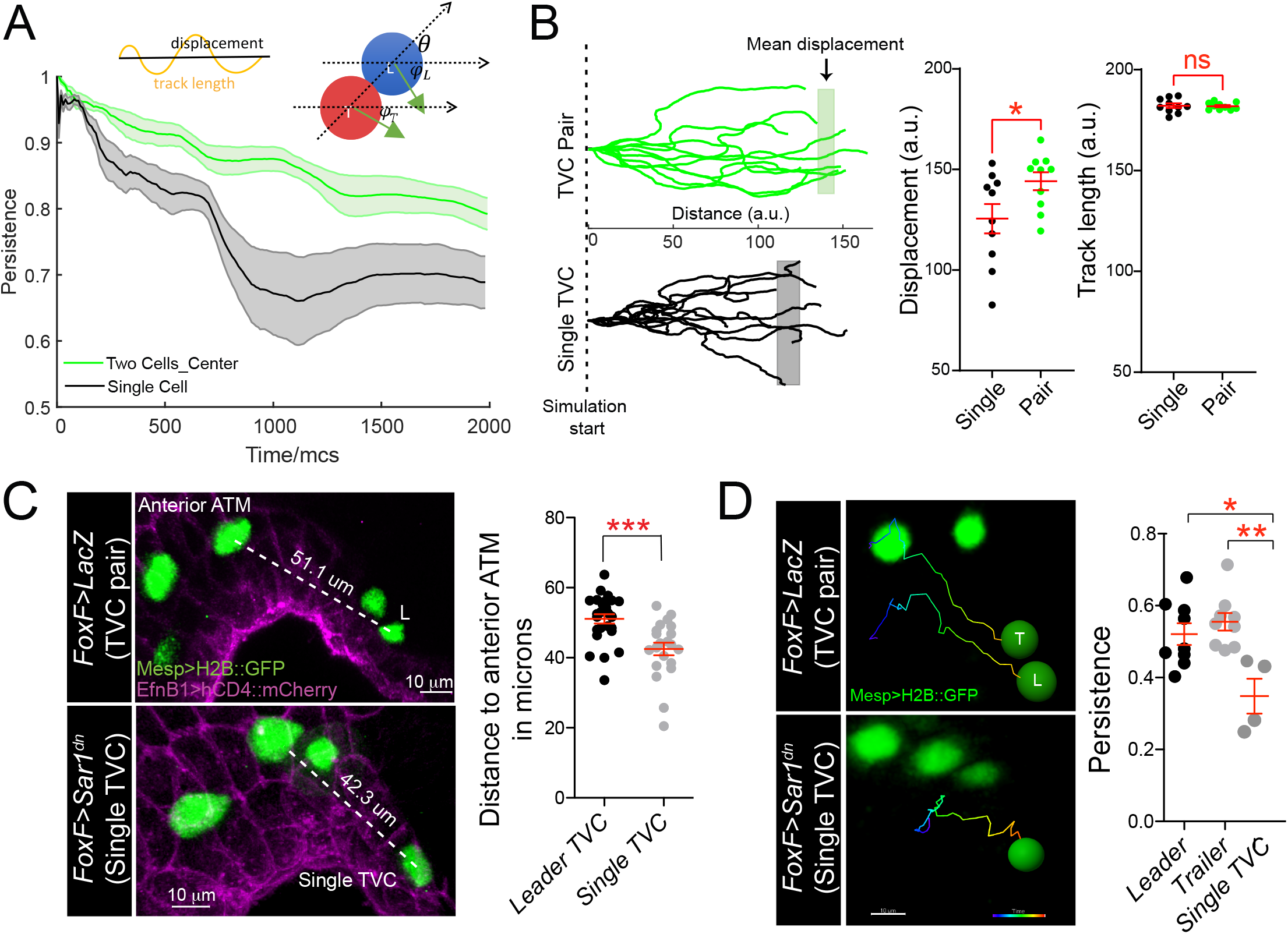
Migratory persistence of cell pairs and single cells. **(A)** Left: Simulation of migration persistence over time for single cell and the centroid of the cell pair. The shaded area shows the standard error. In the model green arrows show the direction of active forces for each cell which is directionally biased but also fluctuates randomly. The leader cell is biased to the right which is the direction of the external cue and the trailer cell is biased to the leader cell. *φ*_*L*_ and *φ*_*T*_ are the angles between the green arrows and the right direction and *θ* is the angle between the line connecting two cells and the right direction. The specific stochastic equation for *φ* is in the Methods section using the same angle notations. The diagram on top shows the relationship between displacement and track length and 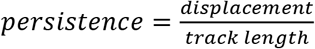. **(B)** Comparisonof tracks from simulations of either migrating cell pairs or single cells within the same simulation time. Shaded vertical lines represent mean final displacement. Graphs on the right show mean total displacement and mean total track length with standard error. Statistical analysis performed using Student t-test. * = p<0.05. **(C)** nvivo analysis of total displacement of leader cells in a cell pair and single TVCs from the anterior ATM. TVC and ATM nuclei are marked with *Mesp>H2B::GFP*, epidermal cell membranes are marked with *EphB1>hCD4::mCherry*. Scatter plot shows average displacement and standard error. *** = p<0.001. **(D)** Invivo migration of TVC pairs compared to single TVC. Nuclei of the cells are used to track cell migration path in 4D data sets. Paths are color coded from early (blue) to late (red). Scatter plot shows mean persistence of leader, trailer, and single TVC with standard error. Statistical analysis performed using 1way ANOVA with Bonferroni post test. * = p<0.05, ** = p<0.01.

Simulation of single migrating TVCs showed these cells to have a smaller overall displacement at the end of migration (Fig. 4B). We assayed the final displacement of single TVCs in vivo to the total displacement of the leader TVC. In agreement with simulations, TVC pairs migrated further away from the anterior ATM than single cells in embryos^30^ (Fig. 4C). To test if this was due to the loss of persistence, as predicted by the simulations above we analyzed the migration path of cell pairs by tracking the nuclei of leader and trailer cells in 4D data sets and compared it to the migration path of single TVCs (Fig. 4D, Supplemental Movies 9, 10). In control conditions the leader and trailer migrate at a similar persistence, however, the migration paths of single TVCs are significantly less straight than those of either leader or trailer cells, suggesting that *in vivo,* collective organization confers robust directionality to the migrating cells.

In summary, both simulations and *in vivo* observations suggested that polarized cell pairs migrate with increased robustness to fluctuations in directionality compared to single cells.

### Polarized cell pairs overcome mechanical resistance of the endoderm during migration

The above sections indicate that collective organization endows the TVCs with defined properties (e.g. persistence) that are intrinsic to the cell pairs and determine the characteristics of their migration. However, TVCs migrate surrounded by other embryonic tissues that are shown to canalize their behavior^29–31^. Specifically, shortly after the onset of migration, the TVCs penetrate the extracellular space between the ventral trunk epidermis, which they use as stiff substrate, and the softer trunk endoderm, which deforms as TVCs progress anteriorly^30^ (Fig. 5A). We reasoned that the trunk endoderm may exert mechanical resistance to the passage of the TVCs, and hypothesize that this resistance may be better overcome by polarized pairs of cells (Fig. 5A, B). While entering the extracellular space, migrating cells presumably experience mechanical resistance from the deforming endoderm. We predicted that, when two cells migrate in a leader-trailer arrangement, the leader exposes to mechanical resistance a surface that is equivalent to that of a single cell, while the trailer pushes from the rear, therefore adding forward-bearing compression force to overcome endoderm resistance (Fig. 5B).

**Figure 5.**
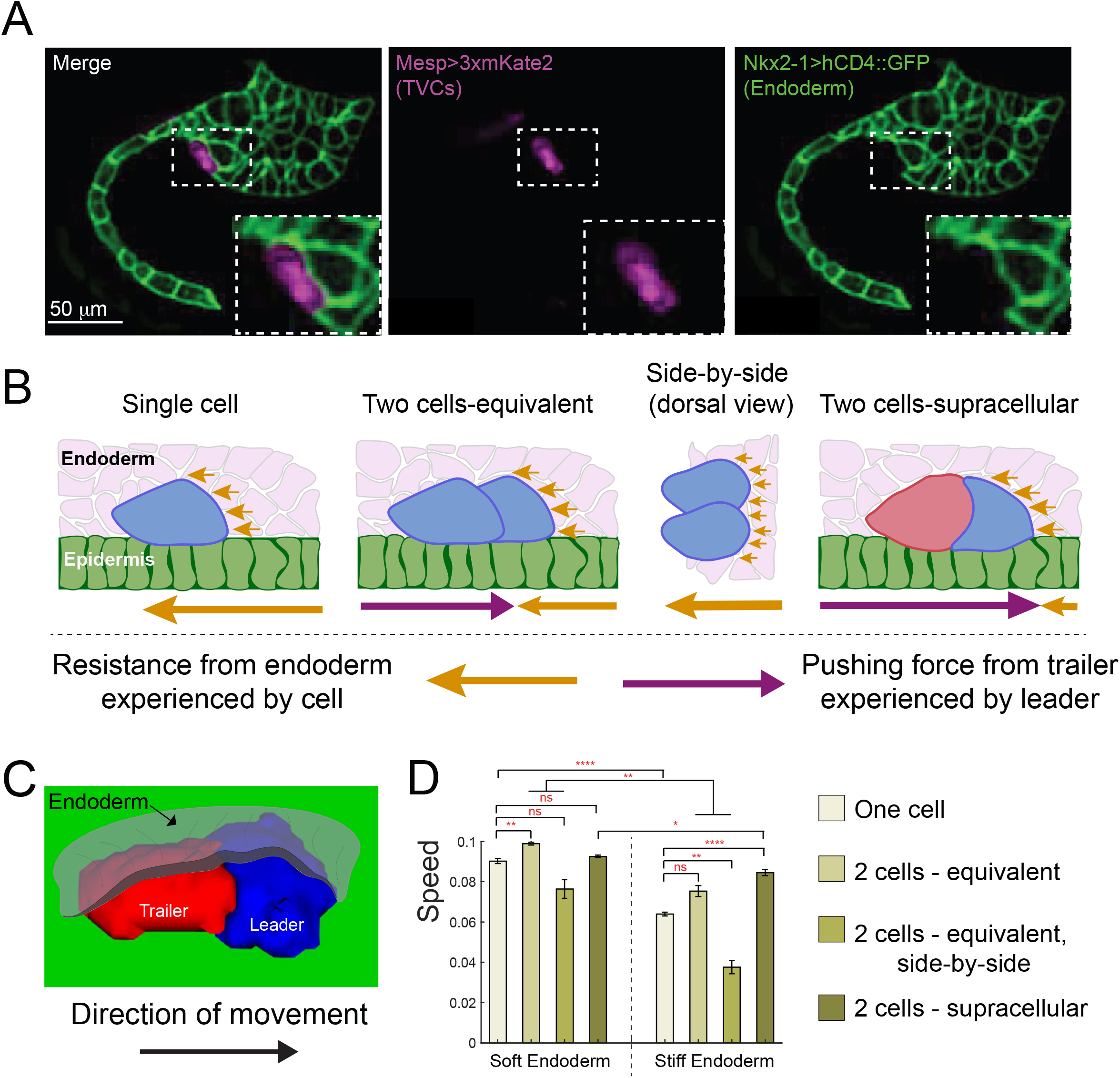
Supracellular cell pairs are more efficient at dispersing pressure from surrounding tissues. **(A)** Micrographs of Stage 23 embryos showing the endodermal pocket formed during TVC migration. Embryos are oriented with anterior to the right. Endodermal cells are marked with *Nkx2-1>hCD4::GFP* (green), TVCs are marked with *Mesp>3xmKate2* (magenta). **(B)** Proposed model for higher efficiency of supracellular cell pairs in overcoming resistance from the endodermal tissue (pink) during migration: the adhesive cell pair shares the resistance force (yellow arrows), which otherwise each single cell must overcome alone. Pressure from the posterior trailer (purple arrows) can help the cell pair overcome resistance from the endoderm. Size of the arrows below the graphics represents relative strength of the force experienced by the cell in the direction of the arrow. **(C)** Simulated supracellular cell pair underneath the endoderm. Epidermis is shown in green. The endoderm is rendered transparent. **(D)** Speed comparison between single cell and differently arranged cell pairs with different profiles and force distributions under the endoderm of varying stiffness. 5 simulations are run for each condition; the error bar is the standard error. Statistical analysis is performed using Brown-Forsythe and Welch ANOVA test. * = p<0.05, ** = p<0.01, **** = p<0.0001

To explore this argument, we simulated cells exposed to varying endoderm stiffness and assayed the effects on migration speed. First, we added deformable endoderm cells to our 2-cell model, and found that the simulated shapes were typical of the supracellular organization (Fig. 5C). Then, we compare speeds of 1- and 2-cell systems by tracking the simulated cells’ centers-of-mass when migrating under an endoderm, with 5 simulations run per condition shown in Fig. 5D). When modeling a soft endoderm, polarized pairs of cells, performed best, whether in supracellular mode or equivalent force distributions, while side-by-side cells were slower, presumably because they expose a greater surface to mechanical resistance (Fig. 5B,D). When simulating a stiffer endoderm, the advantage of supracellular organization became apparent. In these simulations, the supracellular collective migrated faster than all other arrangements (Fig. 5D). Notably, cells migrating side-by-side were slower than single cells with either endoderm stiffness (Fig. 5D), which is consistent the notion that a more extended surface of contact with the endoderm exposes them to greater mechanical resistance, while the smooth teardrop-like shape of single cells may be near optimal for lowering the resistive deformations of the endoderm. Finally, we noted that pairs of equivalent cells migrated slightly faster than the supracellular system under a soft endoderm. By contrast, the supracellular system was the fastest under the stiff endoderm (Fig. 5D). This suggests that the supracellular organization is optimal for minimizing the mechanical resistance of the surrounding tissue to collective migration.

The predicted relative speeds of cells migrating under the endoderm are based on the simplest model, which assumes that the endoderm’s primary effect is mechanical resistance to deformation. *In vivo*, the interactions between TVCs and endoderm cells are likely more complex^31^. In summary, these combined *in silico* simulations and *in vivo* observations indicate that the collective organization of migrating cardiopharyngeal progenitors allows them to overcome mechanical resistance to deforming the endoderm, and reach a typically mesodermal position in between germ layers for prospective cardiac organogenesis.

## Discussion

Complex multicellular behaviors, including directed collective cell migration, emerge from the context-specific integration of universal and dynamic processes, which operate at subcellular scale and are coordinated within and across cells^65^. The sheer complexity of integrated cellular systems constrains direct experimental interrogations, but mathematical models and simulations provide a powerful complement to probe the relative biophysical contributions of defined subcellular processes to cellular behavior.

In this study, we used a mathematical model, built from first biophysical principles, to generate computational simulations and explore the morphodynamic space of motile pairs of cardiopharyngeal progenitor cells in the tunicate Ciona. Qualitative comparisons with experimental data first indicated that the defined shape of cell pairs, similar to that of a single motile cell, emerges from the distribution of higher protrusive activity and cell-matrix adhesion to the leader cell, whereas the rear of the trailer cell is the primary site of myosin-based retraction. The latter prediction was corroborated by *in situ* patterns of myosin activity. This illustrates that a purely mechanical model such as CPM, which assumes that most active stresses are generated at the cellular periphery in addition to hydrostatic pressure in the cytoplasm, can uncover the biomechanical underpinnings of collective cell shape and movement.

The above patterns of protrusive activity, cell-matrix adhesion and contractility might seem trivial, considering the well-established organization of individual migrating cells. However, in the “supracell”, the distribution of various cytoskeletal activities across all cells in a collective suggests the existence of mechanisms to ensure such division of labor. This simple prediction implies manifold roles for the cell-cell contact, in addition to its anticipated low surface tension evoked above.

First and foremost, cell-cell adhesion must exert enough force orthogonal to the junction plane to maintain the integrity of the pair and permit mechanical coupling, lest cells lose contact and migrate disjointly (Supplemental Fig 1C). Conversely, the model predicts that excessive cell-cell adhesion would antagonize cell-matrix adhesion and disrupt collective polarity. Balancing cell-cell and cell-matrix adhesion may result from either mechanical interaction, as suggested by the model, and/or biochemical cross-talks, as observed in other systems^66, 67^. The coexistence and contributions of both cell-cell and cell-matrix adhesion to supracellular migration emphasized the hybrid nature of such multicellular systems, where cells adopt intermediate states on an epithelial-to-mesenchymal continuum^11, 65, 68^.

Close cellular contacts probably facilitate the propagation of direct mechanical and biochemical interactions that underlie supracellular migration. We tentatively distinguish “information flows” that propagate in either a back-to-front or a front-to-back fashion^12, 13^. For instance, similar to *Xenopus* cranial neural crest cells, Ciona cardiopharyngeal progenitors appear to focus contractility at the back of the trailer cell, thus suggesting that a “rear-wheel” engine may help power their migration. However, by contrast with neural crest cells and *Drosophila* border cells, contractility itself is not organized in a supracellular fashion and there is no cell flow within the “cluster”. This “rear-wheel” drive represents a back-to-front mechanical input, which propagates as a compression force and emerges from rear-localized myosin activity, possibly in response to chemorepulsive inputs integrated by the trailer. It is also conceivable that cell-cell adhesion complexes suppress myosin-based contractility at the back of the leader cell, for example through recruitment of Rho GAP molecules by cadherin, as is the case in early *C. elegans* embryos^69^. In neural crest cells, contact-mediated inhibition of locomotion (CIL) provides such front-to-back signals that polarize the cell collective, in part through Cadherins, Ephrin receptors and Planar Cell Polarity (PCP) pathway molecules^13^. The PCP pathway offers a particularly tantalizing explanation for the spontaneous alignment of three or four adhering cells following ectopic induction of the cardiopharyngeal progenitor fate, and the existence of a “leader-trailer” mode of collective arrangement predicted by the model.

TVCs’ collective polarity is marked by higher protrusive activity and cell-matrix adhesion in the leader, as indicated by experimental observations and model predictions^29^. Mechanically, it is likely that adhesion complexes are established following lamellipodia formation and support traction forces, which complement rear-driven compression to propel the cells forward. It is conceivable that lower protrusive activity in the trailer limits the deployment of cell-matrix adhesion complexes. However, cell-matrix adhesion in the trailer is probably needed to anchor the cell and allow for its hydrostatic pressure to push the leader. Therefore, one must invoke (1) mechanisms whereby the leader suppresses protrusive activity in the trailer, while permitting the establishment of trailer cell-matrix adhesion in its path. In Drosophila border cells leader-driven suppression of protrusive activity in follower cells is mediated by Rac^16^, and by Delta-Notch signaling between tip and stalk cells during angiogenesis^70^. It is thus likely that similar “front-to-back” mechanisms govern the collective distribution of protrusions, and by extension cell-matrix adhesion complexes, in cardiopharyngeal progenitors.

While future work combining biophysical modeling, force measurements and/or inference from quantified cell shapes is needed to elucidate the mechanisms underlying supracellular organization *in vivo*, our model and experimental investigations uncovered important consequences for directed migration: namely persistence and mechanical interaction with surrounding tissues. Specifically, both computational simulations and live imaging indicated that the instantaneous directionality of single cells fluctuates more than that of cell pairs. In other words, polarized cells pairs were more persistent. It is possible that cell pairs are better at buffering the noise inherent to navigating a complex and changing environment, in part by distributing interactions over greater surfaces, and integrating guidance cues more accurately.

Finally, our observations indicate that supracellular organization determines the outcome of interactions with surrounding tissues during migration. We previously determined that TVC pairs migrate onto the extracellular matrix (ECM) associated with the basal lamina of the ventral trunk epidermis^31^ which presumably offers a stiff substrate permitting traction forces. Of note, a specific collagen, *col9-a1*, secreted from the trunk endoderm is deposited onto this ECM and necessary for TVC-matrix adhesion and collective polarity^31^. Here, we found that the trunk endoderm resists deformation by migrating TVCs, which can nonetheless move forward by aligning and joining forces to push against and deform endodermal cells to penetrate the extracellular space. Our combined simulations and experimental observations thus suggest a remarkable effect of supracellular organization on the inter-tissue balance of forces that determine morphogenesis in the embryo.

## RESOURCE AVAILABILITY

### Lead Contact

Further information and requests for resources and reagents should be directed to and will be fulfilled by the Lead Contact: Yelena Bernadskaya.

### Material Availability

This study did not generate new unique reagents.

### Data and Code Availability

The code generated during this study is available on GitHub (https://github.com/HaicenYue/3D-simulation-of-TVCs.git)

## EXPERIMENTAL MODEL AND SUBJECT DETAILS

Wild caught *Ciona robusta* (formerly *Ciona intestinalis* type A) were purchased from Marine Research and Educational Products (MREP, San Diego). As invertebrate chordates, animal care approval was not needed. Prior to use animals are housed in a recirculating artificial seawater aquarium under constant illumination to prevent spawning.

## METHOD DETAILS

### Electroporation and transgene expression

*Ciona robusta* (formerly known as *Ciona intestinalis* type A) adults were purchased from M-Rep, San Diego, CA. Gamete isolation, fertilization, dechorionation, and embryo incubation were performed as previously published^36, 37^ The amount of DNA electroporated varied from 10 μg to 90 μg. Animals were reared at 22-24°C. Embryos used for direct visualization of fluorescent markers were fixed in 4% MEM-FA for 30 minutes, cleared with an PBST-NH_4_Cl solution (50mM NH_4_Cl, 0.15% Triton-X100, 0.05% Tween-20 in 1x PBS), mounted in 50% glycerol supplemented with 2% Dabco 33-LV antifade reagent (Sigma-Aldrich, #290734) and imaged using a Leica SP8 X Confocal microscope.

### Live imaging and TVC tracking

To generate 4D datasets, embryos at 4.5 hours post fertilization (hpf) FABA stage 15 were mounted on glass bottom microwell Petri dishes (MatTek, part# P35G-1.5-20-C) in artificial seawater. Plates were sealed by piping a border of vaseline and 5% (v/v) mineral oil (Sigma, #M841-100 ml) and covered with a 22×22 Fisherbrand Cover Glass (# 12-541-B). Embryos were imaged on a Leica inverted SP8 X Confocal microscope using the 40x water immersion lens at 512×512 resolution every 3.5 minutes for 4 to 5 hours. B7.5 lineage nuclei and epidermal cell membranes were visualized using *Mesp>H2B::GFP* and *EfnB>hCD4::mCherry*, respectively, and TVC migration was tracked using Bitplane Imaris Software Spots module.

### Drug treatment

Embryos were electroporated with the myosin binding intrabody iMyo-GFP^38^ (Mesp>Sf9Myo::GFP) and a membrane marker, Mesp>hCD4::mCherry. At 6 hpf embryos were treated with the Rock Inhibitor Y-27632 (Millipore Item #688001) with a final concentration of 10 mM in buffered artificial seawater. At 8 hpf (Stage 23) embryos were fixed in an isotonic 4% formaldehyde solution (MEM-FA, 3.7% formaldehyde, 0.1M MOPS pH 7.4, 0.5M NaCl, 2mM MgSO4, 1mM EGTA pH 8) for 30 minutes at room temperature. Embryos were then cleared by washing 3x in a PBST-NH4Cl and mounted on slides as described above. .

### Image acquisition

All images were acquired using the Leica SP8 X WLL Confocal microscope using the 63x glycerol immersion lens, NA = 1.44. Z-stacks of fixed embryos were acquired at the system optimized Z-step, 512×512 resolution, 600 Hz, and bi-directional scanning. Multiple HyD detectors were used to capture images at various wavelengths.

## QUANTIFICATION AND STATISTICAL ANALYSIS

### Morphometrics analysis

The membrane marker Mesp>hCD4::GFP was used to segment the TVCs and derive morphometric measurements such as sphericity, area, and volume in Bitplane Imaris using the Cell function with cell segmentation calculated from cell membranes with an average cell size of 6. Thresholding is adjusted based on individual image properties. Z-steps were normalized to achieve equal voxel size in X, Y, and Z planes. TVCs were then segmented and resulting cells were exported to separate surfaces. To calculate the distance or angle between cells a point was placed at the center of mass for each cell using either the nucleus or the cell object using the Imaris Bitplane Measurement module.

### Image preprocessing

All 3D stacks were imported and converted to Imaris format. Images were reoriented and cropped with the leader cell to the left. An automated Gaussian filter and background subtraction was applied to all images using the Imaris Batch module. Images were projected to 2D using Maximum Intensity Projection and exported as TIFFs for analysis in FIJI.

### Aspect ratio calculation

2D projected images were imported into FIJI and converted to 8-bit format. A threshold was applied to each individual image and empty spaces were filled using the Binary −> Fill Holes function. Resulting object was used to derive the aspect ratio using the Analyze Particles function. In simulation, black-and-white images were obtained using imbinarize function in Matlab and then regionprop.BoundingBox function was used to get the minimal rectangle that enclose the object. Aspect ratio is the ratio between the width and length of this rectangle.

### Myosin intensity analysis

Images were imported into FIJI. Using the freehand line function with a width of 10 units, a scan was performed on the leading edge of the leader cell, the cell-cell junction, and the membrane of the trailer cell, using the membrane marker as a guide. The intensity of iMyoGFP and the membrane marker Mesp>hCD4::mCherry along the scan was measured using the Plot Profile function and exported as intensity along the line scan. This was done for each cell pair. Readings along the line scan were aligned based on the starting position of the scan and averages were calculated.

### Statistical analysis and data representation

For all data comparing two samples of continuous variables the Wilcoxon Rank Sum test (also known as the Mann-Whitney test) is used. Categorical data is analyzed using Fisher’s Exact test. For data sets containing more than two conditions and taking into account cell type (Leader/Trailer) a two-way ANOVA followed by the Bonferroni posttest was used. For all data sets containing nominal variables a chi-square test is used. P values as reported as follows: *=p<0.05, **=p<0.01, ***=p<0.001.

### Model

We use Cellular Potts Model^39^ to simulate the movement of one or several cells on the substrate. The model is based on the minimization of the effective energy *H* which is a function of the cell shape and areas of contact between adjacent cells. It is computationally efficient to study the multiple 3D cells with enough resolution. In our model, the cell size is about 10×10×10 pixels. The model also allows adding protrusive and retractive forces to the cells^33, 40, 41^ .

In Cellular Potts Model, the space is divided into pixels, and each pixel *i* is assigned a spin *σ*_*i*_. The spin is effectively an index that identifies which cell the pixel belongs to. A stochastic modified Metropolis algorithm^42^ is used to determine how the spin *σ* changes. At each step, the algorithm randomly selects a target site, *i*, and a neighboring source site *j*. If they belong to different cells, or to a cell and neighboring the environment, the algorithm sets *σ*_*i*_ = *σ*_*i*_ with probability, 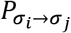, which is determined by the Boltzmann acceptance function:

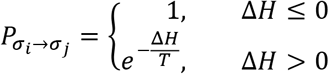

where *ΔH* is the change of the effective energy caused by this change of spin and *T* is an effective temperature parameter describing the amplitude of stochastic fluctuations of the cell boundary^43^. We use *T* = 10 for all the simulations in this paper. The key part of any specific Cellular Potts model is the effective energy *H*. In our model, we define *H* as:

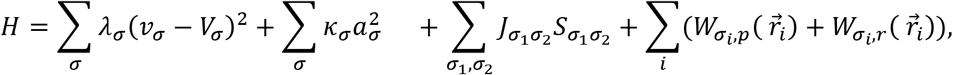

Here the first and the second term represent the effects of the volume conservation and cell surface (cortex) contraction, respectively. Alternatively, the first term can be thought of as the effect of the hydrostatic pressure of the cytoplasm, and the second term – as the effect of the cell cortex tension. The third term represents the adhesion energy between the neighboring cells and between the cells and the extracellular matrix (also called substrate or ECM below).The last term is the effective potential energy related to the protrusive and retractive forces (with subscript *p* and *r* respectively). *i* is the pixel’s index and *σ* is the cell’s or environment’s spin. Variables *ν*_*σ*_ and *a*_*σ*_ are the volume and surface area of the cell *σ*, and *V*_*σ*_ is its target volume. Unlike in some variants of the Cellular Potts Model, we keep the target surface area equal to zero, so effectively the cortex is contractile for any area. The target volume is a parameter that we take to be equal to the cube of the characteristic cell size. Parameters *λ* and *κ* are the coefficient determining how tightly the volume is conserved and how great the cortex tension is, respectively. Parameter 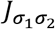 is the adhesion energy per unit area of the boundary between cells *σ*_*1*_ and *σ*_*2*_ (or between cell and environment). Variable 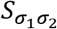 is the area of the boundary between cells or between one cell and the environment. Essentially, the model’s first two terms tend to minimize the cell’s area while keeping its volume constant shaping the cell into a sphere. Adhesion terms, however, try to maximize the boundary areas, flattening the cells. The competition between these terms make individual cell look like a dome on the substrate (this is how we choose relative strengths of the cortex tension and characteristic adhesion), and two cells – like two domes pressed into each other side-by-side. To make the cells move, we must add the forces pushing the cell front and pulling its rear. Note that those forces originate from the cytoskeleton inside the cells, and not in the environment surrounding the cells, so the force balances are implied. Specifically, the force of protrusion that pushes on the cell leading surface forward from inside is balanced by a reactive cytoskeletal pushing directed to the rear and applied to the firm adhesions between the ventral surface of the cell and extracellular matrix. Similarly, the force of retraction that pulls the cell rear forward from inside is also balanced by a reactive cytoskeletal pulling directed to the rear and applied to the firm adhesions between the ventral surface of the cell and extracellular matrix.

We introduce these forces through effective potential energies as follows. First, we define a polarity for each cell, which is quantified using the angle between the polarization direction and the positive-x direction, *φ*, as shown in Fig. 4A. Then, the respective potential energies can be defined as:

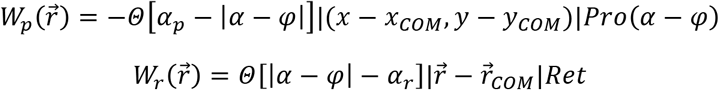

Here *Θ* is the Heaviside step function (equal to 1/0 for positive/negative values of argument, respectively).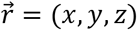 is the 3D position of a specific pixel. *α* is the angle of vector (*x* − *x*_*COM*_, *y* − *y*_*COM*_), in which (*x*, *y*) is the position of 2D projection of a specific pixel onto the x-y-plane, and (*x*_*COM*_, *y*_*COM*_) is the 2D position of the centroid of the cell. *φ* is the polarity angle mentioned above. *α*_*p*_ and *α*_*r*_ are the angular ranges of the protrusive and retractive forces, respectively. For example, if the protrusive force exists in the front half of the cell and the retractive force exists in the back half of the cell, 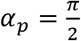, 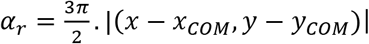 is the 2D distance in x-y plane between specific pixel and the centroid of the cell and by taking gradient of it, we will get the protrusive force in the radial direction in x-y plane which is parallel to the substrate. 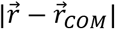 is the 3D distance between a specific pixel and the centroid of the cell and taking a gradient of it, results in the centripetal retractive force. Similarly, taking a gradient of the protrusive part of the effective energy results in the protrusive force which is parallel to the substrate and splays out radially in the xy-plane. In the SI, we explain why the protrusive force are chosen to be parallel to the substrate, while retractive forces are centripetal. *PrO*(*α* − *φ*) and *Ret* define the amplitudes of the energy terms which are also strengths of the protrusive and retractive forces, respectively. *Ret* is a constant parameter, while *PrO* is constant in some simulations but is a function of angle (*α* − *φ*) in others. Their values, as well as the values for *α*_*p*_ and *α*_*r*_, and for all other model parameters, are listed in the Table in the Supplemental Material.

When investigating directionality and persistence of the cells’ trajectories, stochasticity is introduced to the polarity’s dynamics as follows:

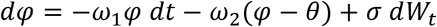

where *θ* is the angle shown in Fig. 3B and *dW*_*t*_ denotes a Wiener process (stochastic directional noise). The first term shows the tendency of the polarity to align with the external signal’s direction (the positive-x direction) and the second term shows the tendency to follow the other cell. For different polarization modes, *ω*_1_ and *ω*_2_ take different values. More specifically, for the Independent mode and the Faster-Slower mode shown in Fig. 3B (when the cells follow the environmental directional guidance independently), *ω*_1_ ≠ 0, *ω*_2_ = 0 for both cells, while for the Leader-Trailer mode (when the trailing cell follows the leader instead of following the environmental guidance), *ω*_1_ ≠ 0, *ω*_2_ = 0 for the leader and *ω*_1_ = 0, *ω*_2_ ≠ 0 for the trailer.

It is worth mentioning that the exact absolute values of parameters in the energy function are not important, as the dynamics of the system is determined by the ratio 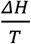 in which *T* is a “temperature” parameter without direct relation to the biological processes, and the “Monte Carlo Step” in the simulation is not directly related to an actual time scale. So, we only check whether the ratios of the model parameters are consistent with the experimentally estimated orders of magnitude of the biophysical parameters.

Experimental estimates of the force generated over 1*μm* of the lamellipodial leading edge are ~1000*pN* and the total traction force exerted by the cell is ~10^4^ − 10^5^*pN* ^30, 44^. As the leading edge of the lamellipodia is only ~0.1 − 0.*2μm* thick, while in our model we cannot generate very thin protrusions, we distribute the total forces generated by the lamellipodia almost uniformly to the whole front of the cell and use the protrusive force density ~100*pN*/*μm*^2^ assuming the height of the cell is ~10*μm* (Fig. 1A). Similarly, we distribute the total traction force uniformly to the back of the cell resulting in the retractive force density ~10^2^ − 10^3^ *pN*/*μm*^2^. Thus, the orders of magnitude of the protrusive and retractive forces are close, and we keep them close in the model. The energy of adhesion between the cell and the substrate is estimated as follows. Each integrin attachment complex has a force ~10 − 30*pN* ^45^ associated with it. The size of an integrin-based adhesion complexes formed at cell contacts with the ECM is ~1 *μm*^2^,^46^ so we estimate the adhesion force densities as ~10 − 30 *pN*/*μm*^2^. Then, the ratio between the active forces and the adhesion forces is ~10. In our model, the force strength parameters, *PrO*(*α* − *φ*) and Ret are on the order of 10 to 100 in dimensionless units, and the adhesion strength parameter *J* ranges from 0 to 20 in dimensionless units, which is consistent with the force ratios from the experimental measurements.

After the orders of magnitude of the protrusion, retraction and adhesion energies are chosen as described, the rest of the principal model parameters are chosen following the following logic. The cortex contractility parameter *κ* is chosen so that an individual non-motile cell has a shape close to that of a hemi-sphere; if this parameter is too small, the cell becomes a ‘pancake’, if too large – a ‘ball’. The parameter regulating the tightness of the cell volume control, *λ*, is fine-tuned to avoid: 1) freezing the cell shape – when this parameter has a value that is too great, most fluctuations of the cell shape get arrested, and 2) loosening the cell shape too much – when this parameter has a value that is too small, cell transiently becomes too small or too large which disagrees with the observations. The values of the parameters for the stochastic directionality experiment are chosen so that the persistence of the single cell predicted trajectories fit that of the observed trajectories. Finally, note that the parameters for the adhesion strength (J) are scaled as follows. We make this parameter a large positive number for the boundary between the two motile cells and endoderm (or for cell-free space boundary in simulations without endoderm); this corresponds to the ‘no adhesion’ regime. Then, the adhesion parameters for cell-cell and cell-ECM boundaries are smaller positive numbers. Thus, the energy in the system decreases when the relative areas of the cell-cell and cell-ECM boundaries increases, so those are the adhesive surfaces.

Note that the dynamics of the simulated cell is determined by the probability function of spin changes, which is defined, by the exponential function with a cut-off at 1. This leads to a speed-force relation as shown in Fig.S3 which is of an exponential form when the force is too small and is saturated when the force is too large. We avoid this artifact because the parameters we choose restrict our simulations to the regime where the speed-force relation is approximately linear.

Simulations were done using software CompuCell3D 3.7.8 ^43^. In the Supplemental Table, we list all model parameters that are varied between different simulations, and in the SI we explain the reasons for the variance. When we simulate the actively migrating cells in the presence of the endoderm that mechanically resists the deformations, we make the endodermal cells mechanically more passive than the migrating cells (their contractile tension is half that of the migrating cells), and vary the endodermal ‘tightness of the volume conservation’ parameter *λ* a few fold less and greater than that of the migrating cells, respectively).

## KEY RESOURCES TABLE

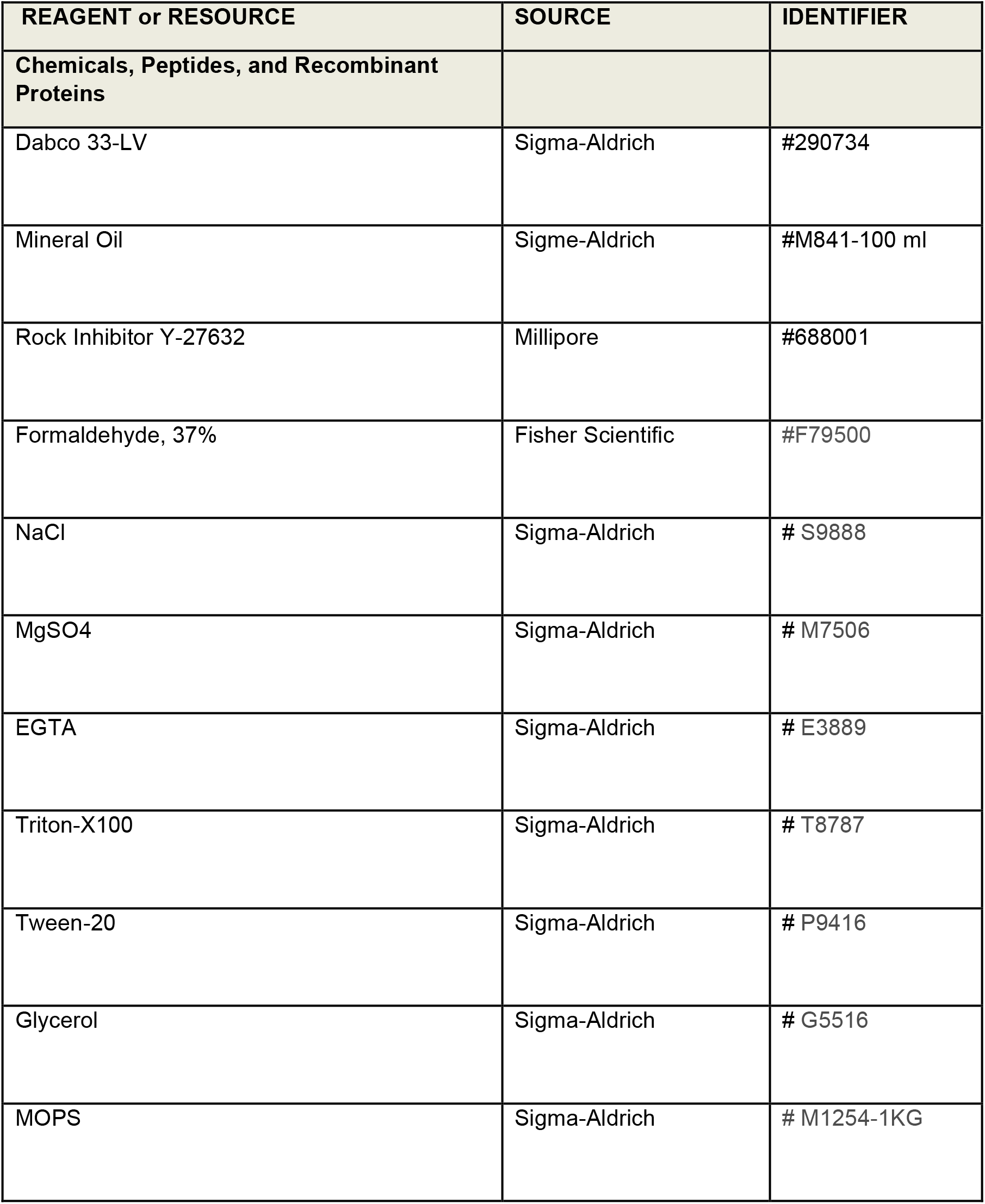
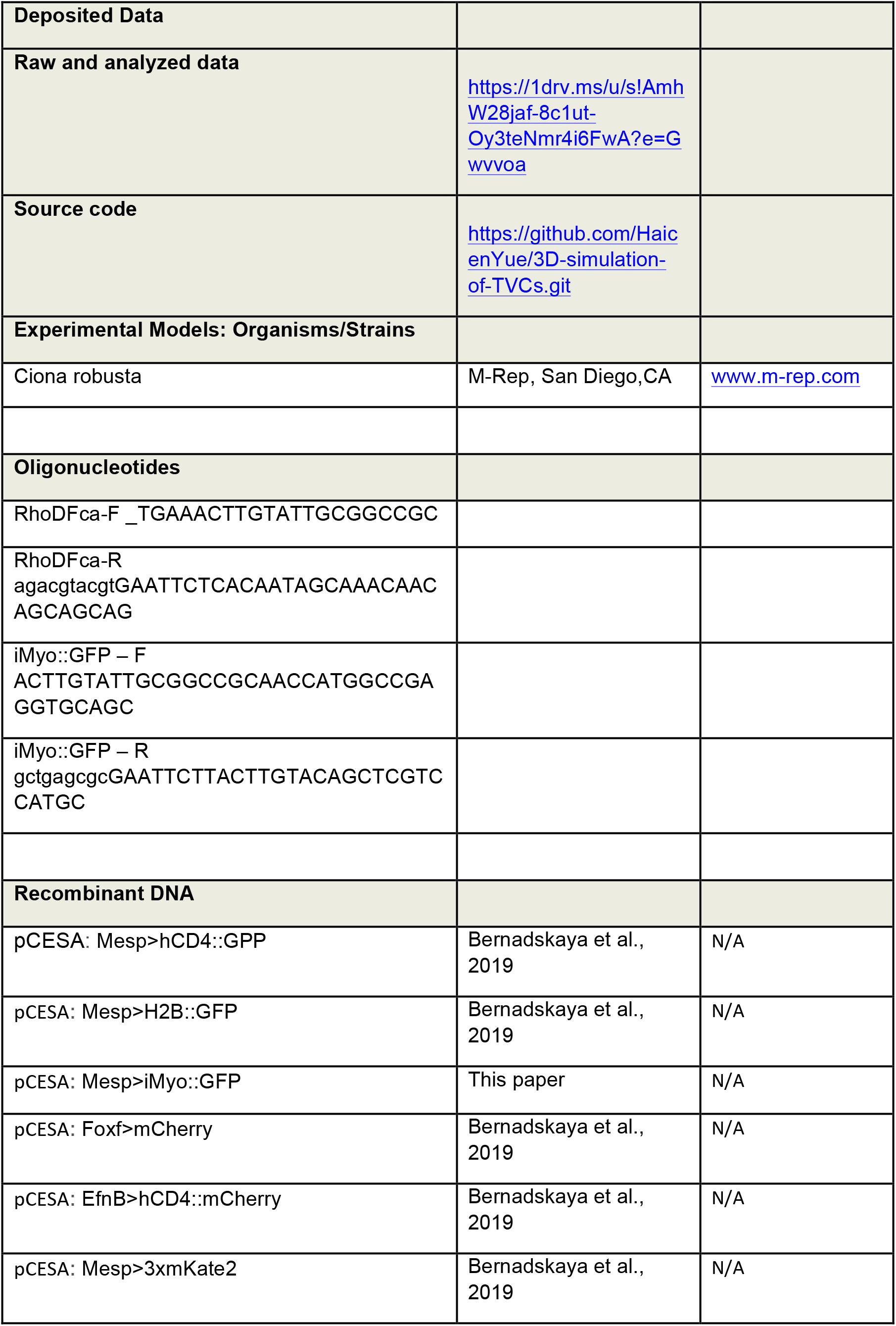
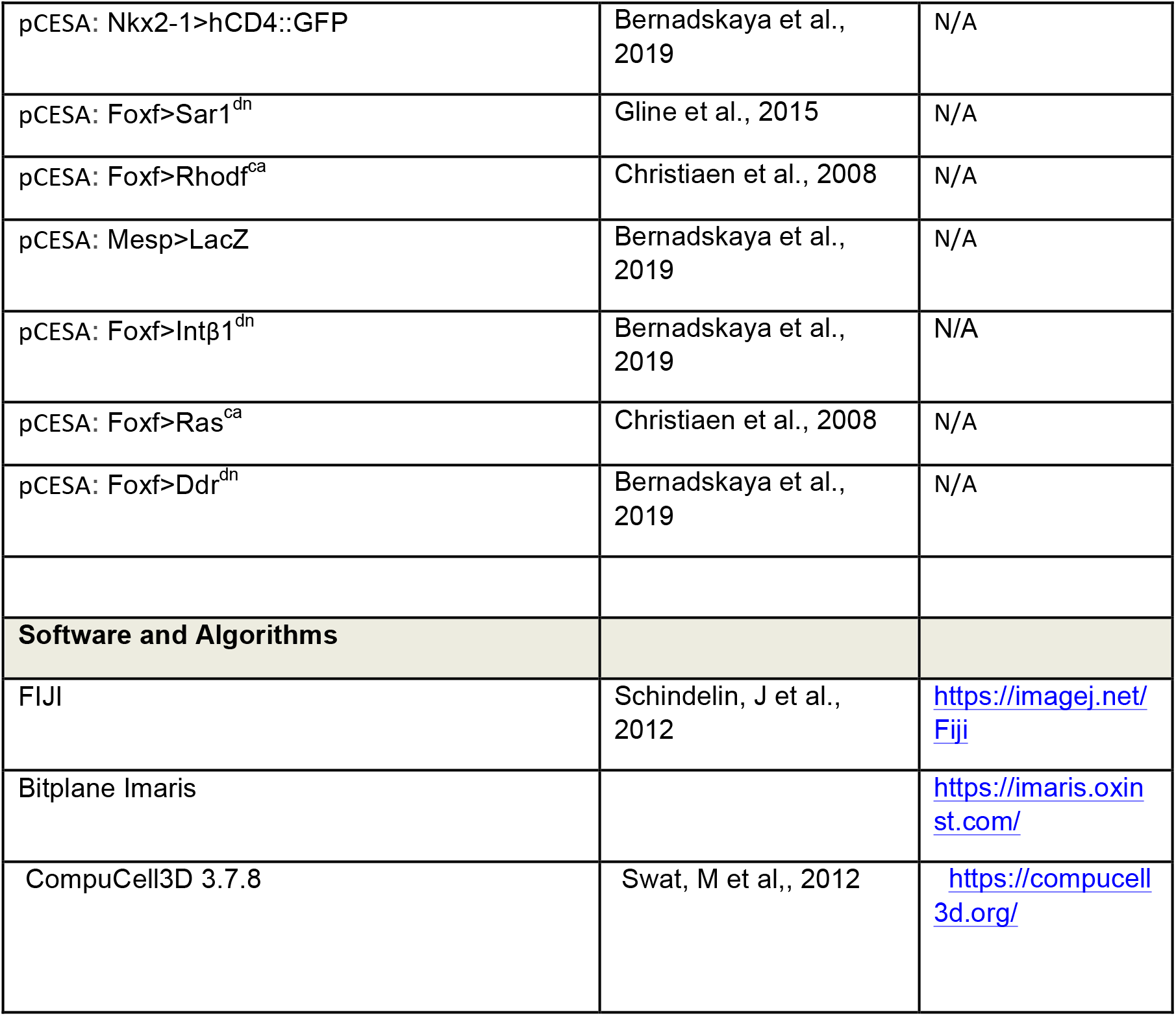

## Acknowledgements

We thank Alex McDougall for his generous gift of the iMyo reporter. This work was supported by NIH F32 GM108369-01A1 post-doctoral fellowship to Y.Y.B, NIH/NIGMS GM096032-09 award to L.C., and NSF #1953430 to A.M.

## Author Contributions

Conceptualization, Y.Y.B., H.Y., C.C., L.C., A.M.; Methodology, Y.Y.B., H.Y., L.C., A.M.; Investigation, Y.Y.B., H.Y.; Writing, Y.Y.B., L.C., A.M.; Funding Acquisition, Y.Y.B., L.C., A.M.

## Supplemental Information

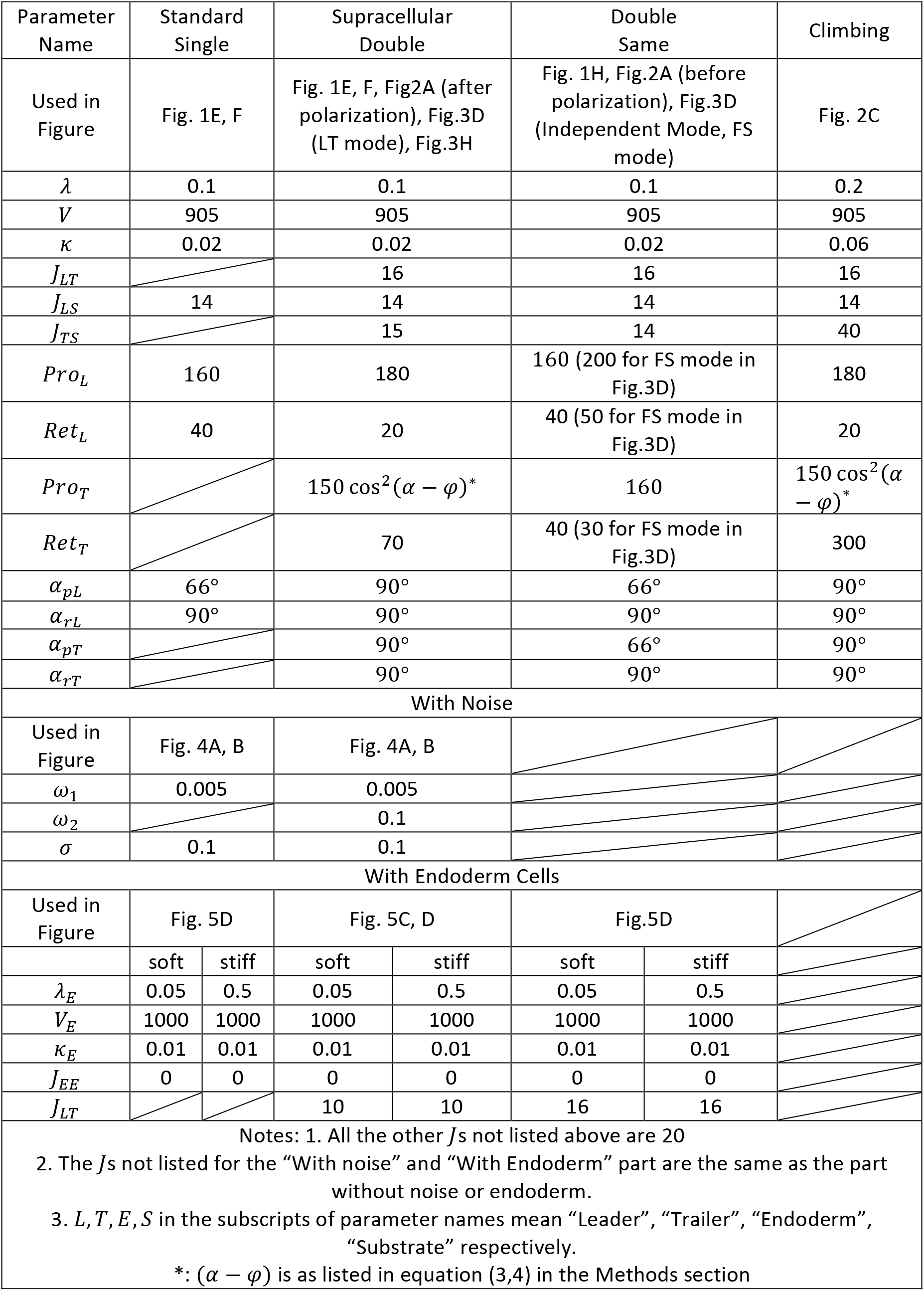
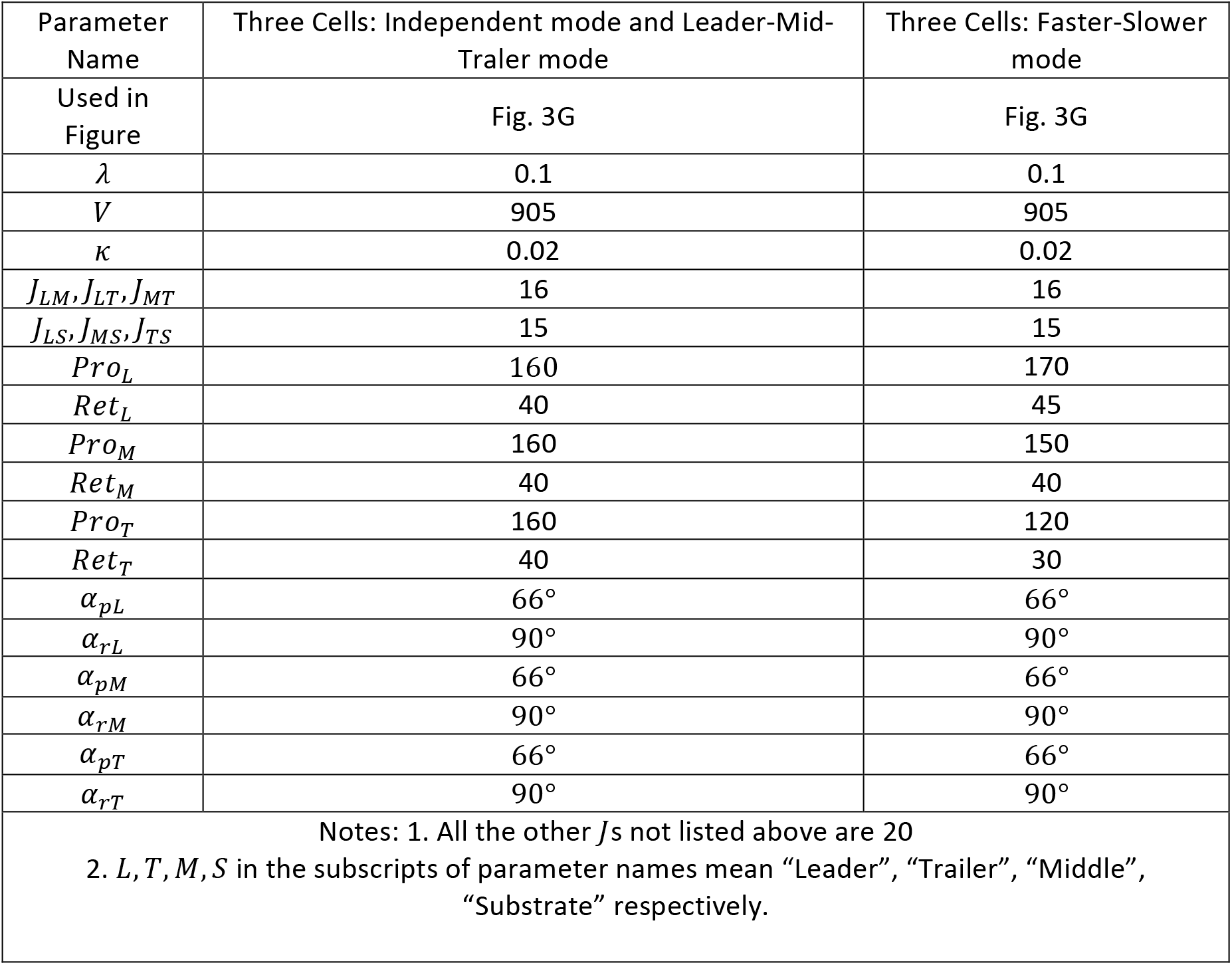

### Arguments for choosing specific spatial-angular force distributions

#### Single Cell

Before we determine the force distribution for the TVC cell pair, we first study the force distribution in a single cell. As shown in Fig. S1A, if the retractive force is concentrated narrowly at the very rear of the cell (if viewed from above), then the cell shape deviates from that observed. Therefore, we chose to distribute the retractive force spatially over the whole rear half of the cell. Two simplest choices for orientation of the retractive forces are: 1) parallel to the substrate, 2) centripetal (the force is inward, along the radial lines from the cell centroid). We decided in favor of the second option, because: a) this option recapitulates the observed tapered cell rear with better than the parallel-to-the-surface force, and b) the origin of this force is likely stress fibers connecting the rear dorsal and central ventral surfaces of the cell, and this geometry is closer to the second option.

One attractive hypothesis would be that the protrusive force is generated by actin filament polymerization of the thin lamellipodial network at the ventral surface of the cell. However, if we restrict the protrusive force only near the bottom of the cell, we must increase the magnitude of the protrusive force by a few-fold in order to maintain its total contribution. After a simulation with such protrusive restricted to the bottom 1 or 2 pixels, with a magnitude equaling 400, the shape of the cell become wavy and unstable, as shown in Fig. S1B. Thus, we relax this restriction and assume that the protrusive force is parallel to the substrate and distributed uniformly in the z direction in the front half of the cell. In principle, the protrusive force could be oriented centrifugally in the x-z plane, but we leave exploration of such option for the future. This choice, of course, raises the question of how such forces are generated. One possibility is that the directional actin network protrudes parallel to the substrate from the firm cell cortex along the whole height of the cell. Future experiments will have to address this question. Then, we need to determine the distribution of the forces in the x-y plane. In principle, the protrusive forces in the x-y plane can be parallel to the x-axis. However, if the magnitude of such force does not depend on the y coordinate, we observed in the simulation that the cell leading edge and side become flat and too wide. Therefore, to avoid introducing an additional parameter to grade the force, we chose the radial centrifugal distribution of the protrusive force in the x-y plane, which reproduces the characteristic leading edge shape without an additional parameter.

In Fig. S1A, we show the typical cell shapes for protrusive and retractive forces distributed within different ranges. We find that decreasing the range of retractive force or increasing the range of protrusive force makes the cell wider and oppositely, the cell becomes longer. Based on the aspect ratio, we finally decide that the retractive force is distributed uniformly in x-y plane in the whole back half of the cell while the protrusive force is distributed uniformly in x-y plane within the range of angle *α* = 66° as shown in Fig. 1D.

#### TVC cell pair

TVC cell pair is not a simple combination of two independent cells. We have shown in Fig. 1H that for two independent same cells moving together, the leading edge of the trailer cell is too wide, which suggests the protrusive force is too widely distributed, and the polarity of the cell pair is not well maintained. This implies that when two cells move together, their force distribution changes because of the cell-cell interactions. So, we decrease the protrusive force in the trailer cell and make it more concentrated along the central axis of the cell by multiplying the magnitude of the force by cos! *α*. This is also consistent with the observation in vivo that in most cases, the two cells’ junction is concave if we look at it from the trailer cell. In addition, we need to increase the retractive force in the trailer cell accordingly to maintain the total force approximately the same so that the speed of the trailer cell does not decrease a lot. This is also reasonable biologically as the generation of protrusive and retractive force share some same cytoskeletal components. We also show in Fig. S1C that if we do not increase the retractive force accordingly, the trailer cell moves slower and will detach from the leader cell. (We observed in general that when the sum of the protrusive and retractive force in the trailer deviates too much from that in the leader, the cells split apart.) We can also observe in vivo that the shape of the leader cell is quite different from a single cell (Fig. 1E). The TVC pair as a whole has a similar aspect ratio as a single cell (Fig. 1F) while the leader cell itself becomes wide and short. This implies that, contrary to the trailer cell, in the leader cell the contribution of the protrusive force increases while the contribution of the retractive force decreases. So, we decrease the magnitude of the retractive force in the leader cell and increase that of the protrusive force while at the same time make it more widely distributed in the cell by setting *α* = 90°. We also show in Fig. S1D that if we keep the force distribution in the leader cell the same as that in the single cell, the leader cell is too long and thus the cell pair looks more like two independent cells connected rather than a supracellular pair, with the aspect ratio similar to a single cell. This is the train of arguments for determining the force distribution in a cell pair based on that in a single cell.

Our main idea is that when two cells are moving as a cell pair, they redistribute their forces by decreasing the force near the cell-cell junction while increasing the forces at the front and back of the pair. But the forces near the junction cannot be decreased too much. We show in Fig. S1E that if the retractive force in the leader and the protrusive force in the trailer are decreased to zero, we will have a leader cell with a long tail as there is not enough force to push the cell-cell junction forward.

It is worth mentioning that the model predicts qualitatively reasonable shapes and movements not only for the chosen set of parameters; varying most of the parameters a few-fold still gives reasonable results. Variation of the model parameters from one numerical experiment to another is shown in the Supplementary Table; mostly, the logic of such variation should be clear from the description of the numerical experiment. Just two notes about the cases when this logic may not be so transparent: when we simulate the trailer ‘climbing’ the ‘leader’, not only the adhesion of the trailer to the substrate is decreased, but also the tightness of the volume conservation and the surface tension are increased, otherwise the cell at the bottom becomes too flat. When we simulate cells migrating under the endoderm, the cell-cell adhesion is adjusted so that the characteristic supracellular shape is conserved.

**Supplemental Figure 1.**
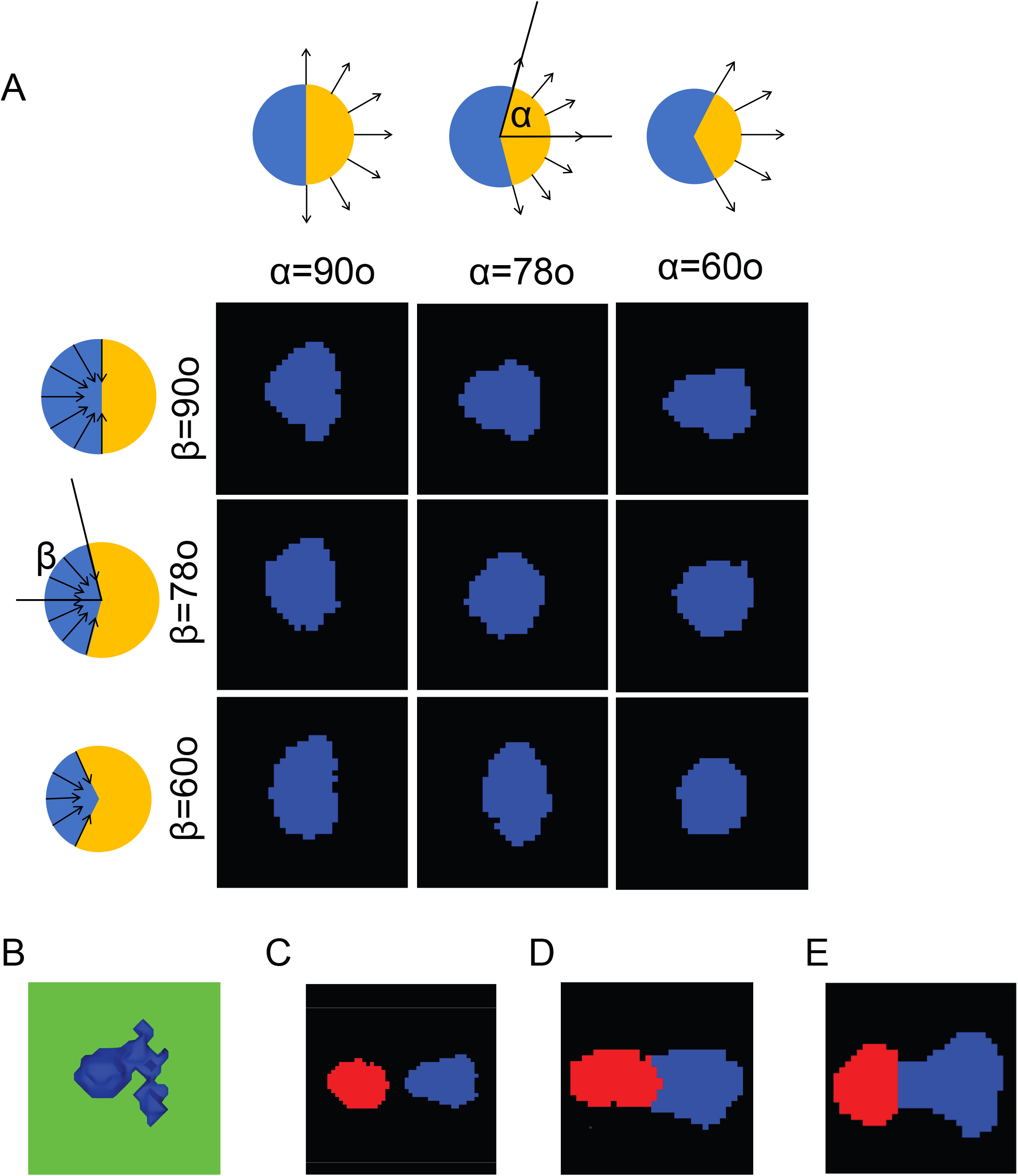

